# Developmental and MAPK-responsive transcription factors drive distinct malignant subtypes and genetic dependencies in pancreatic cancer

**DOI:** 10.1101/2020.10.27.357269

**Authors:** Pasquale Laise, Mikko Turunen, Alvaro Curiel Garcia, Lorenzo Tomassoni, H. Carlo Maurer, Ela Elyada, Bernhard Schmierer, Jeremy Worley, Jordan Kesner, Xiangtian Tan, Ester Calvo Fernandez, Kelly Wong, Urszula N Wasko, Somnath Tagore, Alexander L. E. Wang, Sabrina Ge, Alina C. Iuga, Aaron Griffin, Winston Wong, Gulam A. Manji, Mariano J. Alvarez, Faiyaz Notta, David A. Tuveson, Kenneth P. Olive, Andrea Califano

**Author notes:** These authors equally contributed to this work (Pasquale Laise, Mikko Turunen). These authors equally contributed to this work (Alvaro Curiel Garcia, Lorenzo Tomassoni).

## Abstract

Despite extensive efforts, reproducible assessment of pancreatic ductal adenocarcinoma (PDA) heterogeneity and plasticity at the single cell level remains elusive. Systematic, network-based analysis of regulatory protein activity in single cells identified three PDA Developmental Lineages (PDLs), coexisting in virtually all tumors, whose transcriptional states are mechanistically driven by aberrant activation of Master Regulator (MR) proteins associated with gastrointestinal lineages (GLS state), morphogen and EMT pathways (MOS state), and acinar-to-ductal metaplasia (ALS state), respectively. Each PDL is further subdivided into sub-states characterized by low vs. high MAPK pathway activity. This taxonomy was remarkably conserved across multiple cohorts, cell lines, and PDX models, and harmonized with bulk profile analyses. Cross-state plasticity and MR essentiality was confirmed by barcode-based lineage tracing and CRISPR/Cas9 assays, respectively, while MR ectopic expression induced PDL transdifferentiation. Together these data provide a mechanistic foundation for PDA heterogeneity and a roadmap for targeting PDA cellular subtypes.

## Introduction

Pancreatic ductal adenocarcinoma (PDA) is the third-leading cause of cancer-related mortality and is highly resistant to cytotoxic, targeted, and immune therapies (Rahib et al., 2014). Compared to the heterogeneous mutational repertoire of other cancers, PDA is remarkable for its relatively uniform complement of DNA alterations, with frequent mutations in *KRAS*, *CDKN2A*, *TP53*, and *SMAD4*. Unfortunately, these hallmark events are not yet targeted by approved therapies and additional mutations known to confer specific drugs are uncommon. Consequently, cytotoxic combinations remain the standard of care, with most patients quickly exhibiting primary or secondary chemoresistance.

Cellular heterogeneity has emerged as a major contributor to cancer chemoresistance, due to potential coexistence of malignant subpopulations with distinct transcriptional states (*i.e., subtypes*) and equally distinct drug sensitivities (Neftel et al., 2019), as well as to the contribution of diverse stromal subpopulations (Elyada et al., 2019). It is thus reasonable to expect that, as observed in other tumors, chemoresistant states may provide effective reservoirs whose plasticity will eventually regenerate the full tumor heterogeneity (Neftel et al., 2019), thus frustrating efforts to delineate therapeutic vulnerabilities through bulk tissue analyses. Stromal cells further complicate matters as they often represent the dominant compartment in bulk PDA samples.

Multiple studies, in large PDA cohorts, agree on the presence of at least two transcriptional PDA subtypes, with more differentiated tumors—corresponding to *Classical* or *Progenitor* subtypes in prior studies—showing association with better outcome, compared to poorly differentiated ones— termed *Quasi-mesenchymal*, *Basal-like*, or *Squamous* (Bailey et al., 2016; Collisson et al., 2011; Maurer et al., 2019; Moffitt et al., 2015; Network., 2017; Puleo et al., 2018). However, the molecular signatures of these subtypes were derived from a mixture of both tumor and stroma related transcriptional states, which are not consistent across individual tumors and datasets (Birnbaum et al., 2017) and may also average across multiple coexisting malignant states. Indeed, published classifiers present limited overlap when applied across available cohorts (Birnbaum et al., 2017). While removing stromal contributions from expression signatures helps harmonize discrepancies (Maurer et al., 2019) a comprehensive assessment of heterogeneity of malignant cell states in PDA remains elusive and represents an important next step for the field.

To address this challenge, we used metaVIPER (Ding et al., 2018), a single-cell implementation of the extensively validated VIPER (Virtual Inference of Protein activity by Enriched Regulon analysis) algorithm (Alvarez et al., 2016). This regulatory network-based algorithm can be used to accurately quantitate the transcriptional activity of any regulatory protein—such as transcription factors, cofactors, and other proteins that participate in direct regulation of a cell’s transcriptional state—from single cell RNA-Seq (scRNAseq) data; this is accomplished by leveraging the expression of its transcriptional targets as a multiplexed reporter assay. For this study, transcriptional targets were identified using the Algorithm for the Accurate Reconstruction of Cellular Networks (ARACNe) (Basso et al., 2005) (**Figure S1A**). We have previously shown that VIPER–assessed regulatory protein activity effectively overcomes a major limitation of single cell profiles, where ≥ 80% of genes typically fail to produce any reads (“*gene dropout”*), and compares favorably with flow cytometry and other antibody-based single cell assays (Elyada et al., 2019; Laise et al., 2022; Obradovic et al., 2021), without the typical limitations due to availability and optimization of such reagents. VIPER has proven effective in identifying Master Regulator (MR) proteins whose activities drives cell transcriptional state (Alvarez et al., 2016), as confirmed by a comprehensive body of literature, see for instance (Alvarez et al., 2018; Aytes et al., 2014; Carro et al., 2010; Rajbhandari et al., 2018).

VIPER analysis of scRNA-seq profiles from transformed single cells dissociated from a total of 35 PDA patients (scRNA-seq), across three public datasets, reproducibly identified six transcriptionally-distinct cell states. These comprise three developmental cell lineages, each one further subdivided into two sub-states distinguished by either high or low RAS/MAPK effector protein activity. We named the three developmental cell lineages *Gastrointestinal Lineage State* (GLS), *Morphogenic State* (MOS), and *ADM-Like State* (ALS), based on functional associations of their activated proteins with early gastrointestinal identity, epithelial-to-mesenchymal transition, and acinar-to-ductal metaplasia drivers, respectively. Within each developmental lineage state, we denote as M^+^ (MAPK active) or M^-^ (MAPK inactive) the sub-states associated with high or low Raf-MEK-ERK signaling. Barcode-based lineage tracing in PDA cell lines representing either a GLS or MOS state provided clear evidence of widespread, plasticity between the M^+^ and M^−^ states, across both lineages, while MEK inhibition, using multiple pharmacological agents, effectively induced M^+^ → M^−^ transition in multiple cell lines, with potential implication for new therapeutics targeting RAS. The six states identified by our analysis were recapitulated in single cell profiles from multiple human PDA cohorts, cell lines, and PDX models. Furthermore, bulk tumor analysis showed that patients with tumors enriched for MRs activated in MOS state (Morphogenic tumors) presented with poorer prognosis and less differentiated tumors compared to patients with tumor enriched for MRs activated in GLS state (Lineage tumors).

To validate VIPER-predicted subtype-specific candidate MRs (i.e., most aberrantly activated, and inactivated proteins), we performed pooled CRISPR/Cas9-mediated viability screens for Lineage and Morphogenic MRs (which were well represented among cell lines) and confirmed they were enriched in proteins essential for viability in subtype-specific fashion. Finally, to validate the role of top VIPER-inferred MRs in mechanistically determining PDA cell state, we showed that ectopic expression of Lineage MRs in Morphogenic tumors effectively transdifferentiated them to the Lineage-like tumors, both at the bulk and at the single cell level. Analysis of perturbed profiles from these assays confirmed the intra-connected and autoregulated nature of the MR modules controlling the Lineage/Morphogenic transition. A conceptual workflow of our overall approach is depicted in **Figure 1A**.

**Figure 1:**
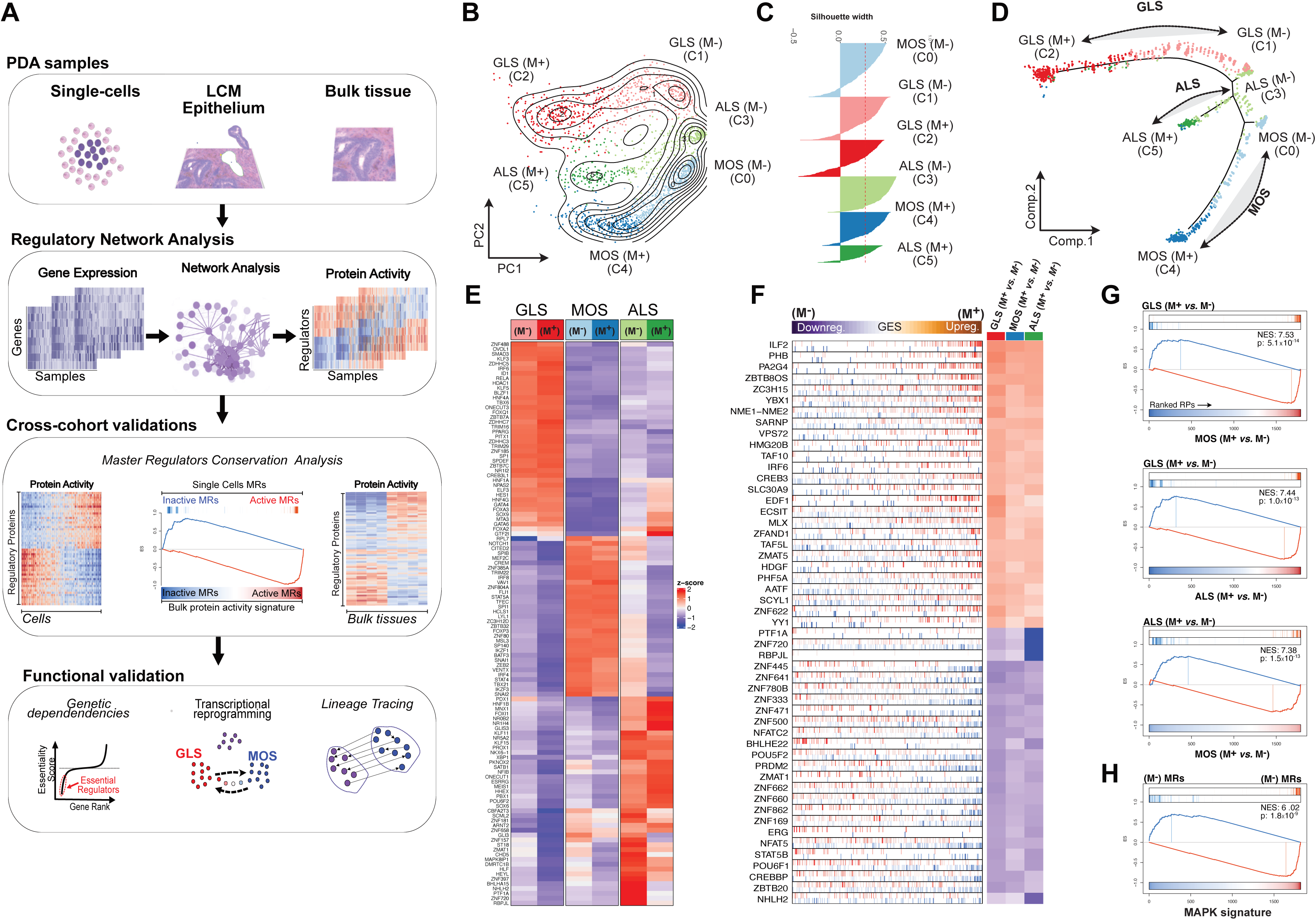
PDA transcriptional cell states. **A)** Conceptual workflow of this work. **B)** Principal Component Aanalys (PCA) based on VIPER-inferred protein activity profiles showing the six cell cell states identified by cluster analysis. **C)** Silhouette plot showing the silhouette score of each individual cell in the assigned cluster. **D)** Pseudo-trajectory plot generated by the Monocle algorithm on the VIPER-inferred protein activity profiles. **E)** Heatmap showing the VIPER-inferred most differentially activated 25 proteins of each cluster compared to all the other clusters, including the GLS and MOS markers (HNF1A, GATA6, NOTCH1, SNAI1, SNAI2,GLI3 GATA4, SOX9,HNF4A). **F)** msViper plot of the MAPK signature identified across PDLs. Each row shows the distribution of the target genes (vertical lines) of a given regulatory protein in the gene expression signature distinguishing M+ vs M-cells (top bar). Only target genes with highest likelihood across the 4 PDA networks used for metaVIPER are shown in the plot. Blue vertical lines represent negatively regulated target genes, red vertical lines represent positively regulated target genes. The top bar represents the integrated gene expression signature across the three PDLs distinguishinng M^+^ vs M^-^ states, with differentially expressed genes ranked from most downregulated in M^-^ (blue) to the most upregulated in the M^+^ (orange). The three columns on the right indicate the activation status of each regulatroy protein in the M^+^ state compared to M^-^ state within each PDL. Red indicates activation in M^+^ state, blue indicates inactivation in the M^+^. **G)** GSEA plots showing the conservation of VIPER-inferred MRs between M^+^ and M^-^ states across the three PDLs. Top, GSEA plot shows the 2-tailed enrichment analysis of the top 50 MRs (hits) (the 25 most activated and 25 most inactivated) distinguishing M^+^ and M^-^ states of the GLS PDL in the protein activity signature (refence signature, x-axis) distinguishing M^+^ and M^-^ states in MOS PDL. Same analysis was done to perfom a pairwise comparisons across all the PDLs. NES and p-value were estimated by 2-tailed enrichment analysis with 1000 permutations. **H)** GSEA plot showing the conservation of the MAPK MRs (the 25 most activated and 25 most inactivated) inferred from patients, in the VIPER MAPK signature experimentally inferred by perturbing Raf-MEK-ERK in PDA cell lines. NES and p-value were estimated by 2-tailed enrichment analysis with 1000 permutations.

## Results

### Regulatory Network Reverse Engineering

To accurately infer PDA-specific transcriptional targets (regulons) for 1835 regulatory proteins (i.e., the PDA regulatory network) from gene expression data, we used the ARACNe algorithm (Basso et al., 2005) (**Table S1**). ARACNe is an information theory-based algorithm with a strong history of experimental validation, see (Califano and Alvarez, 2017) for a review. PDA-specific regulons were independently generated from four cohorts, including the ICGC (Bailey et al., 2016), TCGA (TCGA Research Network: https://www.cancer.gov/tcga), UNC (Moffitt et al., 2015), and CUMC-E. The latter comprises RNA-seq profiles generated from the epithelial compartment of 200 laser microdissected PDA samples collected at Columbia University, representing the expansion of a previously published 68-profile dataset (Maurer et al., 2019). Taken together, the integration of these four networks provides a balanced, consensus-based representation of human PDA, including RNA-seq and microarray-based profiles, varying patient demographics and selection criteria, and differences in stromal infiltration.

### Single Cell Protein Activity Analysis

MetaVIPER was designed to integrate the analysis of multiple networks at the individual protein level (Ding et al., 2018); this is especially useful in single cell analyses, where multiple cell types may co-exist, see for instance (Elyada et al., 2019; Obradovic et al., 2021; Obradovic et al., 2022). We used metaVIPER to measure the activity of the 1,835 regulatory proteins in each of 1856 malignant epithelial cells from six PDA patients using published scRNA-seq profiles (Elyada et al., 2019). Following quality control, malignant cells were affirmed by ploidy (Yuan et al., 2018) and genomic instability analyses, based on inferred-CNVs, (Laise and Alvarez, 2022; Patel et al., 2014) (**Figure S1 B-E**). Regulatory protein activity profiles were then used in all downstream analyses, including cell states identification, cross-cohort analysis, and functional validations assays.

### Identification of Molecularly Distinct PDA Cell States

We have previously shown that protein activity-based cluster analysis is more robust than gene expression-based clustering, including in single cell analyses (Elyada et al., 2019; Obradovic et al., 2021; Paull et al., 2021). Consistently, activity-based Louvain clustering (Stuart et al., 2019) of malignant PDAC cells revealed an optimal solution—based on silhouette analysis (Rousseeuw, 1987)—comprising six molecularly distinct cell states (**Figure 1B-C)** that were not apparent by gene expression analysis (**Figure S1F**).

Protein activity-based pseudotime trajectory analysis, as computed by Monocle (Qiu et al., 2017), produced a branched structure with six transcriptionally distinct states organized into three distinct branches of PDA-specific Developmental Lineages (PDLs) (**Figure 1D**). These predictions were consistent with single cell stratification by principal component analysis (PCA) (**Figure 1B**), showing clear separation of the three PDLs along the second principal component (PC2), with the first principal component (PC1) separating each PDL into two sub-states.

Differential regulatory protein activity analysis was instrumental in characterizing the three PDLs as (a) a *Gastrointestinal Lineage State* (GLS), associated with activation of established gastrointestinal (GI) lineage markers (*e.g.*, GATA4, GATA6, HNF1A, HNF4A, HNF4G) (**Figure 1E**), (b) a *Morphogenic State* (MOS), associated with GI marker inactivation and activation of morphogen pathway and Epithelial to Mesenchymal Transition (EMT) markers (*e.g.*, NOTCH1, GLI3, and ZEB2, SNAI1 SNAI2, respectively), and, finally, (c) an *ADM-Like State* (ALS), associated with activation and overexpression of acinar-to-ductal metaplasia (ADM) markers (*e.g.*, ONECTUT1, SOX9, SPP1) (**Figure 1E and S1G)**. Ploidy and inferred CNV analyses confirmed that ALS cells harbor the same complement of chromosomal copy-number variations as assessed in the other subtypes (Laise and Alvarez, 2022; Patel et al., 2014; Yuan et al., 2018) confirming their *bona fide* malignant nature.

We then focused on characterizing differences between the two molecularly distinct transcriptional states comprising each PDL. This analysis revealed strong conservation of the MR proteins associated with the M^+^ and M^-^ states across all three PDLs (p ≤ 1.5.X10^-13^, by two-tailed GSEA test with 1000 permutations**) (Figure 1F-G)**. Among the most differentially activated MRs, we found TFs associated with RAS signaling, such as YY1 and YBX1, while the most inactivated MRs included PTF1A and RBPJL, established acinar cell regulators that are typically inactivated upon RAS activation (Lin et al., 2020; Yin et al., 2022; Yuan et al., 2017).

Given the dominant role of mutant KRAS in PDA biology, these analyses implied that the two transcriptional substates of each PDL may be related to differential RAS signaling activity. This was supported by the highly significant enrichment of MAPK pathway genes in genes differentially expressed between the M^+^ vs. M^-^ sub-states across all PDLs (p = 3.2 X10^-5^, by GSEA one-tailed test, with 1000 permutations) (**Figure S1H**). To further test this hypothesis, we generated an experimental, PDA-specific MAPK activity signature by integrating the regulatory protein activity signatures of two cell lines (PANC1 and ASPC1, characterized as representative of the MOS and GLS PDLs, respectively, see **Figure S4A**) following treatment with 14 different RAF, MEK, or ERK inhibitors (see methods). As expected, we observed highly significant enrichment of such consensus MAPK activity signature in MRs differentially active in the M^+^ vs. M^-^ sub-state of each PDL, across all samples **(Figure 1H**, p-value= 1.8 X10^-9^, by GSEA two-tailed test, with 1000 permutations). Consistent with the established role of RAS/MAPK signaling in cellular metabolism control, we observed lower expression of numerous metabolic enzymes in the M^-^ vs. M^+^ state of PDA cells **(Figure S1I)**. Finally, we performed single cell analysis on six human PDA cell lines and found that in five of the six, treatment with serum-free media significantly decreased the ratio of cells in M^+^ vs. M^-^ states (**Figure S2A)**. The same effect was observed in an independent experiment in ten out eleven PDA cell lines treated with the MEK inhibitor trametinib **(Figure S2B)**. Taken together, these observations support the naming of these sub-states as either “MAPK-active” (M^+^) or “MAPK-inactive” (M^−^). Thus, each PDA cell state is regulated by a combination of MR proteins controlling either developmental lineage or MAPK activity.

### Cross-Cohort Reproducibility

To assess the reproducibility (i.e., cross-cohort classification consistency) of the six cell states identified in the Elyada dataset (Elyada et al., 2019), we analyzed single cells from two additional, independent cohorts comprising 8,300 and 11,300 transformed epithelial cells, dissociated from 5 (Chan-Seng-Yue et al., 2020) and 24 (Peng et al., 2019) human PDA samples, respectively. The analysis revealed statistically significant enrichment of state-specific MRs in proteins differentially active in single cells from the two additional cohorts, thus confirming high cross-cohort cell state reproducibility, with 98% and 96% of the cells matching the six PDL states from the Elyada set, respectively (**Figure S2 C-E,** *p* ≤ 0.05, Bonferroni-corrected by two-tailed aREA test (Alvarez et al., 2016)). Furthermore, the six PDL states were also confirmed in epithelial cells dissociated from 7 human PDA cell lines (**Figure S2F**) and from a PDX model (**Figure S2G)**. Of note, in contrast to patient-derived samples, virtually all PDA cell lines presented with a majority of the cells in either a GLS (M^+^) or MOS (M^+^) state (**Figure S2F**), while tumor samples from patients presented with a majority of the cells in M^-^ states (**Figure S2D-E**).

### Lineage tracing demonstrates rapid interconversion between the M^+^ and M^−^ states

Pseudo-trajectory analysis predicted a continuum of PDA cells spanning between the M^+^ and M^−^ state in each PDL, suggesting potential plasticity between these states. To address this question, we performed barcode-based lineage-tracing assays (Weinreb et al., 2020) in cell lines representative of the GLS and MOS PDLs (we could not identify cell lines with a sufficient ALS fraction for analysis). For this purpose, we first generated a single-cell reference map of 7 PDA cell lines (**Figure S3A-B**). Among these cells we selected the PATU8988S and KP4 lines for lineage tracing experiment, since they were among the most specifically GLS and MOS cell-state enriched, respectively (**Figure S2F**).

We transduced these two cell lines with an average of ∼3 million random 27-nucleotide barcodes, such that the probability of transducing barcodes with fewer than 4 nucleotide differences in two distinct cells would be vanishingly small (p = 1.4×10^-14^). scRNA-seq profiles from a total of 35,547 PATU8988S and 19,778 KP4 cells were generated, supporting cell fate tracing of 259 KP4 (1.3%) and 512 PATU8988S cells (1.4%), identified by 145 and 325 unique barcodes, respectively (see methods).

Following classification of PATU8988S and KP4 cells into M^+^ and M^-^ states (**Figure 2A-B**, see methods), we assessed barcode representation in the M^+^ and M^−^ states within each cell line, at both an early (*T*_0_ = 6d for KP4 and *T*_0_ = 10d for PATU8988S) and a late (*T*_1_ = 17d for KP4 and *T*_1_ = 38d for PATU8988S, i.e., 10 population doublings) time point following transduction. Among KP4 and PATU8988S cells, 52% and 54% of the barcodes were unequivocally observed in cells that were in two different states at the two time points, indicating plastic interconversion between these states. Specifically, for the PATU8988S line, 106 of 275 M^+^ cells (transduced with 88 of 325 unique barcodes (27%)) spontaneously transdifferentiated to the M^-^ state, while 113 of 237 M^-^ cells (transduced with 89 of 325 unique barcodes (27%)) transdifferentiated to the M^+^ state (**Figure 2C**). For the KP4 line, 78 of 151 M^+^ cells (transduced with 54 of 149 unique barcodes (36%)) spontaneously transdifferentiated to the M^-^ state, while 26 of 108 M^-^ cells (transduced with 24 of 149 unique barcodes (16%)) transdifferentiated to the M^+^ state (**Figure 2D**), see methods. Taken together, these results indicate high M^+^ ↔ M^-^ plasticity in both the GLS and MOS PDLs.

**Figure 2:**
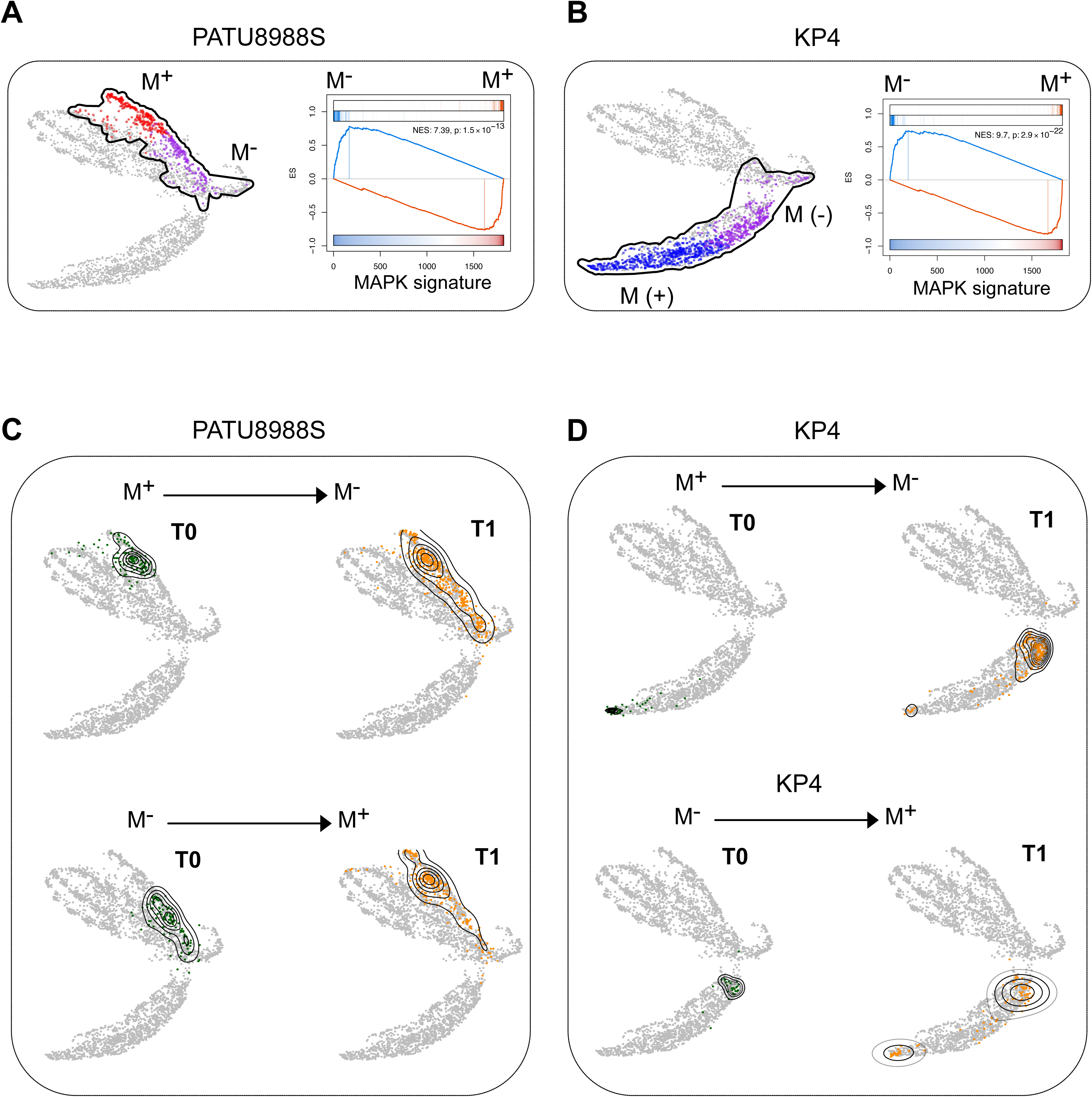
Single-cell lineage tracing of PDA epithelial cells. **A)** Reference map (UMAP) showing the single cells of seven PDA cell lines used as reference (see methods). The black line highlights the area of the reference map enriched for PATU8988S cells. Red dots represent the PATU8988S cells classified as GLS (M^+^) and purple dots represent PATU8988S cells classified as GLS (M^-^). Grey dots represent the cells of all the other cell lines used to generate the refence map. The GSEA plot shows the conservation of the top 50 differentially activated proteins (the 25 most activated and the 25 most inactivated) computed by comparing PATU8988S GLS (M^+^) vs PATU8988S GLS (M^-^) in the experimentally-inferred MAPK VIPER signature in PDA cell lines (reference signature). NES and p-value were estinated by 2-tailed enrichemt analysis with 1000 permutations. **B)** Reference map (UMAP) showing single cells of the seven PDA cell lines used as reference. The black line highlights the area enriched for KP4 cells. Blue dots represent the KP4 cells classified as MOS (M^+^) and purple dots represent KP4 cells classified as MOS (M^-^). Grey dots represent cells of all the other cell lines used to generate the refence map. The GSEA plot shows the conservation of the top 50 differentially activated proteins (the 25 most activated and the 25 most inactivated) computed by comparing KP4 MOS (M^+^) vs KP4 MOS (M^-^) cells in the experimentally-inferred MAPK VIPER signature in PDA cell lines (reference signature). NES and p-value were estinated by two-tailed enrichemt analysis with 1000 permutations. **C)** Top, reference map showing the transition of PATU8988S cells from M^+^ to M^-^ state in the single-cell lineage tracing experiment. Bottom, reference map showing the transition of PATU8988S cells from M^-^ to M^+^ state. **D)** Top, reference map showing the transition of KP4 cells from M^+^ to M^-^ state in the single-cell lineage tracing experiment. Bottom, reference map showing the transition showig the transition of KP4 cells from M^-^ to M^+^ state.

### Differential GLS and MOS Representation Drives Bulk-tissue-based Clustering

We next sought to explore whether cell states identified at the single cell level could recapitulate the subtypes identified by metaVIPER-based bulk PDA sample clustering across four publicly available cohorts—including TCGA, ICGC, UNC, and Collisson et al. Further, to avoid stromal contributions, we also analyzed the epithelial compartment of 200 LCM samples from the CUMC-E cohort. These are richly annotated with survival data, demographic information, clinical variables, and histopathological annotations of adjacent tissue sections (manuscript in preparation). Using metaVIPER to transform each expression profile to regulatory protein activity profiles, we identified two optimal clusters each (using k-medoids) in the well-established ICGC and TCGA cohorts (Bailey et al., 2016)(TCGA Research Network: https://www.cancer.gov/tcga). The MR proteins whose differential activity optimally stratifies the two clusters were nearly identical between the two independently analyzed cohorts (*p* = 10^-40^, by two-tailed aREA analysis, Bonferroni corrected) (**Figure S3C**). Given such remarkable overlap, we used Stouffer’s method (Stouffer, 1949) to integrate the MR activity p-values across the two cohorts to create a single, high-confidence, bulk sample-based MR activity signature (TCGA/ICGC MR signature) (**Figure 3A**).

**Figure 3:**
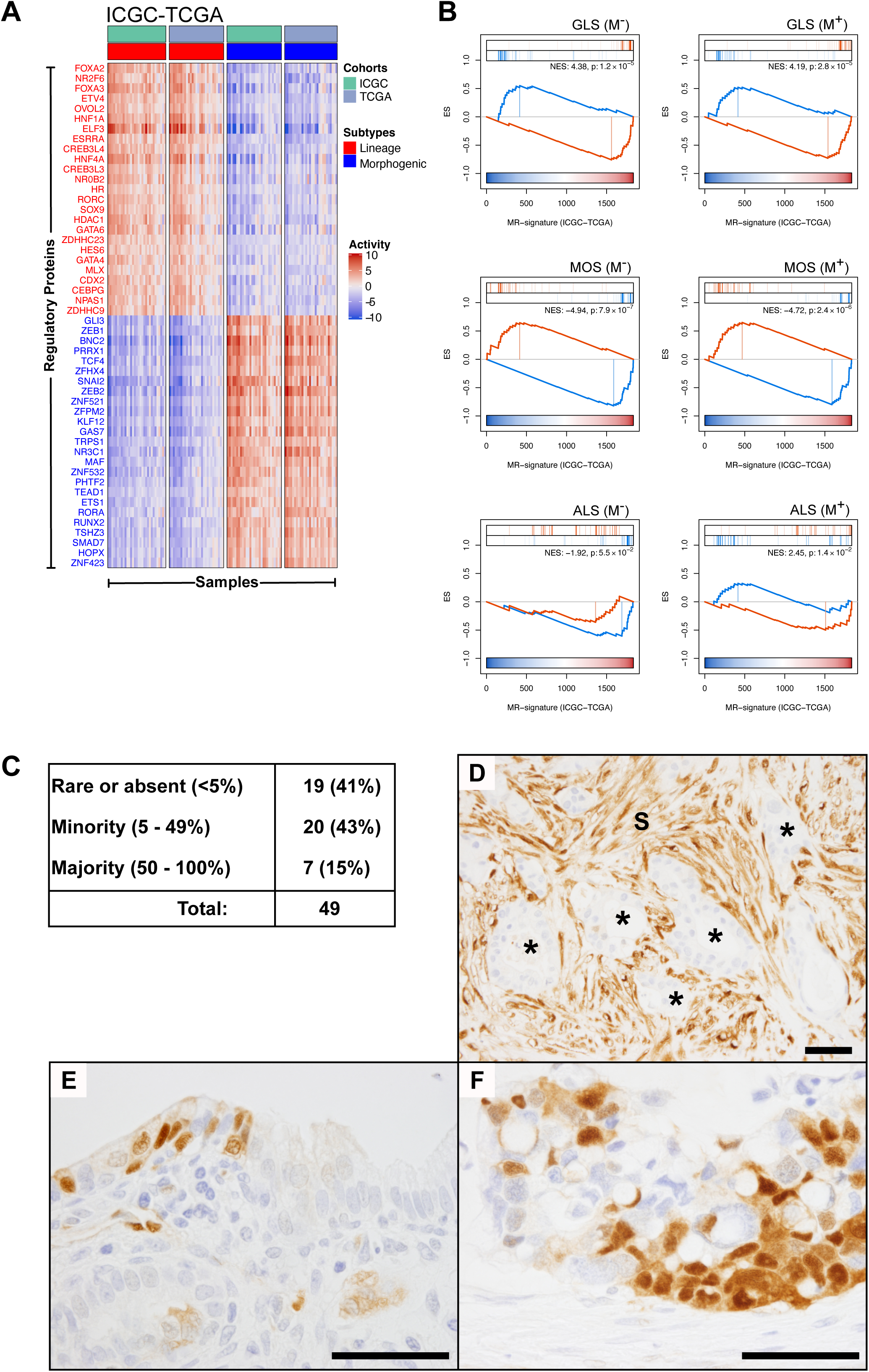
Bulk-based cluster analysis and Immunohistochemistry for phospho-ERK on human PDA cells. **A)** Heatmap showing clusters of TCGA and ICGC patients based on the most differentially active MRs (Tumour Checkpoint) generated by the integration of ICGC and TCGA analyses. One cluster was enriched for GLS MRs and one cluster was enriched for MOS MRs (see figure 3B). **B)** GSEA plots showing the enrichement of the cell state-specific master regulators identified by single-cell analysis in the VIPER-inferred MR signature generated by integrating ICGC and TCGA analyses. NES and p-values were estimated by two tailed enrichement analysis with 1000 permutations. **C)** Table showing blinded scoring of percent pERK positive epithelial cells in a series of 46 human PDA samples from surgical resections. **D)** Example of absent pERK staining in malignant epithelia (*) despite positive stromal cells (S). **E)** Rare positive cells intermixed in a largely negative malignant epithelium. **F)** Malignant epithelium with a majority of pERK positive cells. Scalebars = 50μm.

We then asked whether the results of this analysis recapitulated the cell state MRs identified at the single cell level. MR enrichment analysis confirmed that the clusters from bulk PDA samples closely recapitulated the GLS and MOS single cell states (p ≤ 1.2×10^-5^ and p ≤ 2.4×10^-6^, by two-tailed aREA analysis, respectively) leading to their designation as “Lineage” and “Morphogenic” bulk clusters/subtypes, respectively. We did not observe differential enrichment of MOS and GLS MRs specifically active in M^+^ or M^−^ cell states, probably due to limited molecular resolution of the bulk profiles. ALS state MRs were less enriched in either bulk subtype (p = 1.4×10^-2^). This was expected since the ALS M^-^ state is rather ubiquitously represented in patient derived samples, thus failing to contribute to sample differences, while ALS M^+^ cells are too rare to produce a dominant signature (**Figure 3B**). Indeed, phospho-ERK IHC performed on 48 human PDA samples found that most tumors have very low fractions of active RAS/MAPK signaling within the malignant epithelial compartment, consistent with the observed predominance of M^−^ cells in most human PDA datasets (**Figure 3C-F**). Taken together, these data confirm that bulk sample analyses can only recapitulate two out of the six molecularly distinct subtypes identified at the single cell level.

To further assess reproducibility of the Lineage and Morphogenic clusters identified by the integrated TCGA/ICGC cohort analysis, we assessed whether differential MR activity could be recapitulated in additional cohorts. Indeed, our analyses show that Lineage and Morphogenic MR activity effectively stratified samples in (a) the CUMC-E cohort (**Figure 4A)**, (b) three additional publicly available bulk-level cohorts—including the Moffitt (Moffitt et al., 2015) and Collisson (Collisson et al., 2011) cohorts (**Figure S3D**)—as well as (c) an additional LCM cohort (CSY) (Chan-Seng-Yue et al., 2020) (**Figure S3E)**.

**Figure 4:**
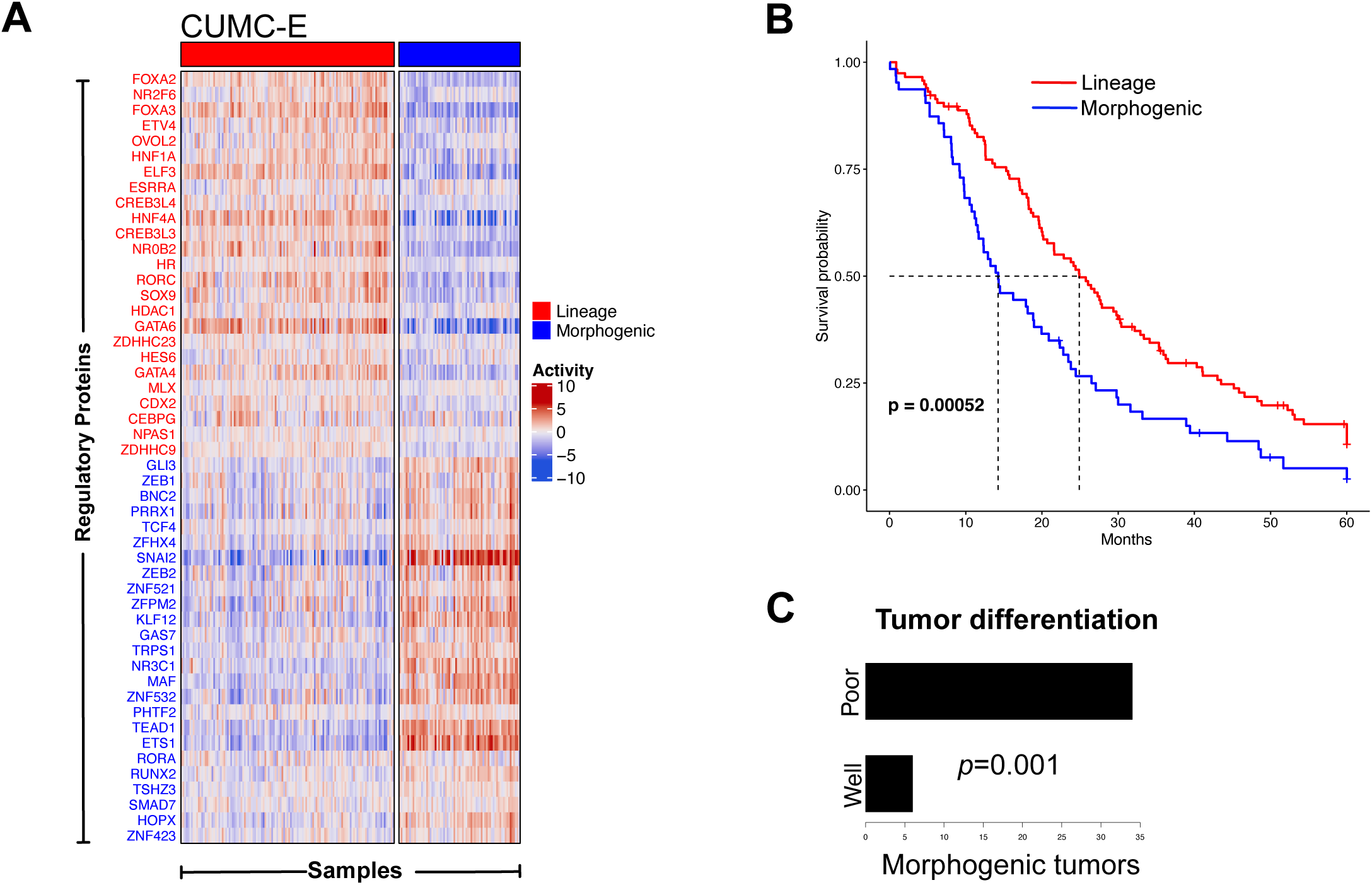
Master Regulator-based classification of CUMC-E cohort. **A)** Heatmap and patients stratification of the CUMC-E cohort based on the MR Tumour Checkpoint derived by the integration of ICGC and TCGA analyses. **B)** Kaplan-meier curve showing that patient stratification of CUMC-E cohort in Morphogenic and Lineage based on MR Tumour Checkpoint correlates with patient’s survival (p-value was computed by log-rank test). **C)** Barplot showing that Morphogenic tumors are enriched for poorly differentiated tumors. The p-value was computed by Chi-square test.

Compared to published gene expression-based analyses, there was significant overlap of the Morphogenic subtype with aggressive subtypes associated with poor survival (*p* <0.05, by one-tailed Fisher Exact Test), including the Quasi-mesenchymal (Collisson et al., 2011), basal-like (Moffitt et al., 2015), Squamous (Bailey et al., 2016) and basal-like A and B (Chan-Seng-Yue et al., 2020) (**Figure S3F**). Consistent with this observation, Morphogenic tumors had significantly worse survival than Lineage tumors in the CUMC-E cohort (*p* = 0.0005) (**Figure 4B**), with a hazard ratio HR = 1.8. Moreover, blind histopathological analysis of adjacent sections from CUMC-E sample blocks showed that Morphogenic tumors were much more likely to be poorly differentiated compared to Lineage tumors (*p* = 1×10^-3^, by *X*^2^ test) (**Figure 4C**). Furthermore, analysis of 190 patients in the CSY set LCM cohort (Chan-Seng-Yue et al., 2020) showed that Morphogenic tumors were significantly enriched for major mutant KRAS imbalance (*p* = 2×10^-4^, by two-tail *Χ*^2^ test) (**Figure S3G**). Finally, while genetic alterations failed to co-segregate with either expression or activity-based subtypes (**Figure S3H**), suggesting an isogenic nature of identified subtypes, epigenetic analysis of TCGA samples revealed a strikingly distinct DNA methylation pattern in Lineage versus Morphogenic tumors (**Figure S3I**). These differences specifically affected key PDL MRs, such as GLS MRs (e.g., GATA6 and HNF1A), which were aberrantly methylated in Morphogenic tumors, and MOS MRs (e.g., ZEB1 and ZNF423), which were aberrantly methylated in Lineage tumors. Taken together these data suggest that bulk cluster-subtypes are mostly driven by differential representation of individual cells in a GLS and MOS states, which are associated with patient outcome, histology, epigenetic state, and KRAS imbalance. Indeed, classification of PDA cell lines in Lineage and Morphogenic based on bulk profiles almost perfectly recapitulates their differential enrichment in MOS and GLS cells at the single cell level (**Figure S4A** and **S2E)**.

### Lineage and Morphogenic MRs represent state-specific dependencies

we then tested whether cell state specific MRs represent critical non-oncogene dependencies essential for cell viability. For this, we focused on the GLS and MOS cell states for two reasons: (a) these states appear to have complete opposite MR activity—i.e., the most activated GLS MRs are among the most inactivated in MOS and vice-versa—and (b) cell lines predominantly representative of the ALS state were not readily available. To assess PDA cell line dependency on state-specific MRs, we performed pooled CRISPR/Cas9 screens in six cell lines—including three recapitulating GLS/Lineage MRs (PATU-8988S, HPAFII, and CAPAN1) and three recapitulating MOS/Morphogenic MRs (PANC1, KP4, and PK45H), using both knock-out (CRISPRko) and inhibition (CRISPRi) systems (**Figure 5A)**. Cells were transduced with a pooled guide-RNA library targeting 3,179 genes (4 sgRNA/gene), including 1835 regulatory proteins (Alvarez et al., 2016) and 759 selected core essential and non-essential genes, as positive and negative controls, respectively (Hart et al., 2017; Palin et al., 2018) (**Figure S4B-C**). Cells were harvested and sequenced at *T*_0_ = 3d and *T*_1_ = 33d, following sgRNA transduction and sgRNA counts were integrated across both technical and biological replicates to assess their differential representation over time and then compared between subtypes to generate a differential, subtype-specific, essential signature. We found that sgRNAs associated with viability reduction in GLS/Lineage and MOS/Morphogenic cell lines were highly enriched in top 200 most differentially active proteins (i.e., 100 most active in GLS and 100 most active in MOS), as produced by metaVIPER analysis of the genes differentially expressed in GLS vs. MOS cells (*p* = 9.1 ×10^-5^, by two-tailed GSEA analysis), as well as in the top 200 MRs obtained by metaVIPER analysis of genes differentially expressed in Lineage vs. Morphogenic samples (p = 2.5×10^-4^) (**Figure S4D**). This was not surprising since GLS and MOS MRs were highly overlapping with Lineage and Morphogenic MRs, respectively (**Figure 3B**). For GLS/Lineage cell lines, the analysis identified CDX2, GATA6 and HNF1A among the most essential MRs, while for MOS/Morphogenic cell lines it identified MYBL1, ZEB1 and GLI2 (**Figure 5B-C)**. The demonstrated genetic dependence of PDAC cell states on state-specific MR proteins offers a potential path for the future identification of drugs that selectively target individual cell types in PDA.

**Figure 5:**
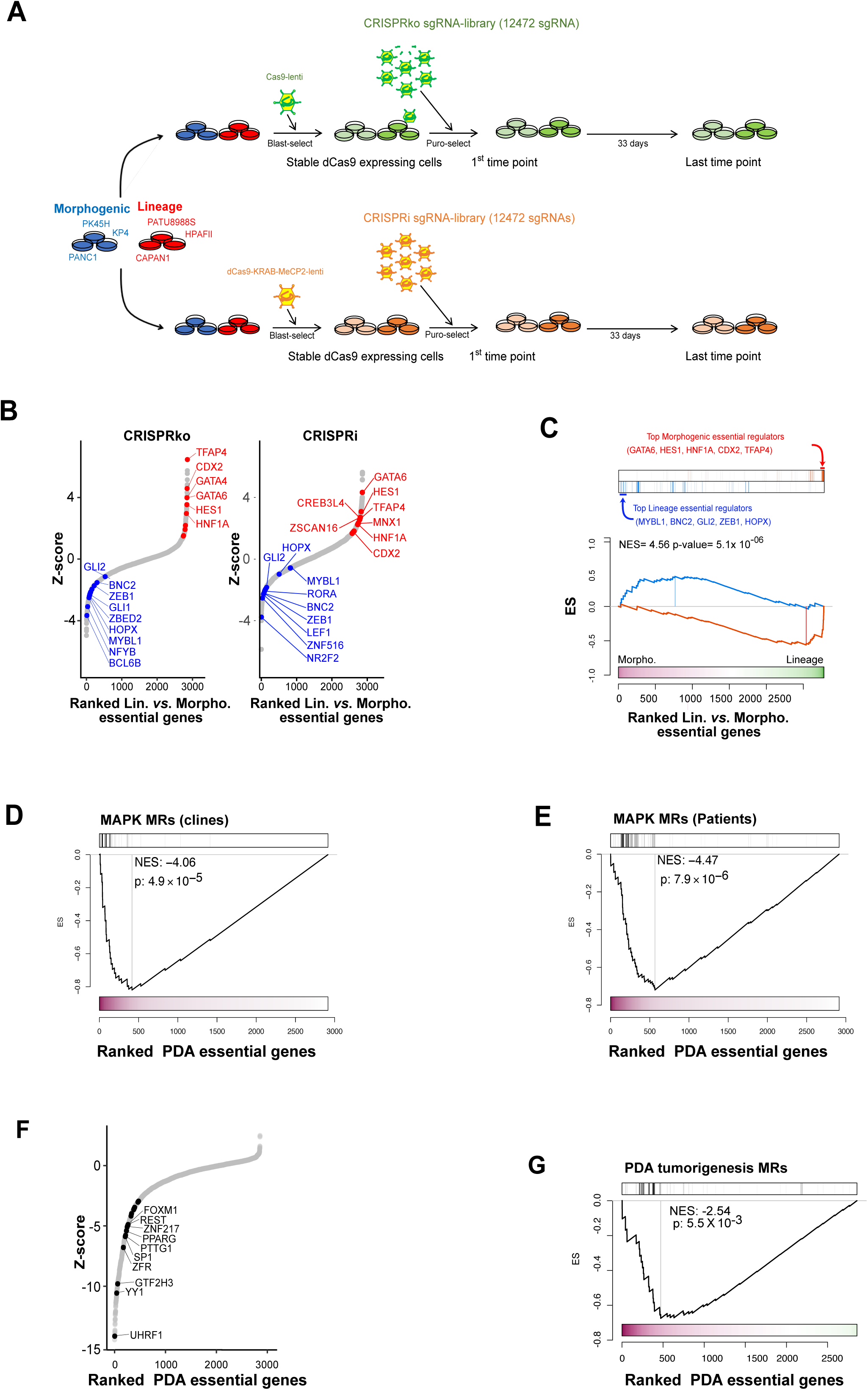
PDA Master Regulators are enriched in essential genes. **A)** Schematic workflow used for the pooled CRISRP/dCas9 (CRISPRi and CRISPRko) screens. **B)** Scatter plots showing differential gene essentiality signature between Lineage and Morphogenic cell lines (Lineage/Morphogenic _ess_), as produced by CRISPRko (left) and CRISPRi (right). Genes are ranked according to their differential essentiality score (z-score), from the most Morphogenic-essential (left) to the most Lineage-essential (right). VIPER-inferred Lineage and Morphogenic MRs are shown in red and blue, respectively. **C)** GSEA analysis shows statistically significant enrichment of the 50 most differentially active Lineage and Morphogenic MRs in genes assessed as differentially essential between Morphogenic and Lineage cell lines, by integration of CRISPRko and CRIPSRi data (p-value and NES estimated by GSEA 2-tailed test, with 1,000 permutations). **D**) GSEA analyses showing statistically significant enrichment of the 50 most differentially active MAPK MRs experimentally inferred by perturbation in PDA cell lines (clines) in the subtype-independent essentiality signatature as determined by integrating CRISPRko and CRISPRi signatures across all the cell lines (p-value and NES estimated by GSEA 1-tailed test, with 1,000 permutations). **E)** GSEA analyses showing statistically significant enrichment of the 50 most differentially active MAPK MRs inferred by single-cell analysis of PDA patients in the subtype-independent essentiality signatature as determined by integrating CRISPRko and CRISPRi signatures across all the cell lines (p-value and NES estimated by GSEA 1-tailed test, with 1,000 permutations). **F)** Scatter plot ranking genes based on their PDA subtype-independent essentiality as determined by CRISPRko, with VIPER-inferred, subtype-independent Master Regulators highlighted. **G)** GSEA of PDA MRs subtype-independent, computed by comparing the 200 LCM CUMC-E samples against the gene expression centroid all normal tissue samples in GTEx (Consortium, 2013), in subtype-independent essential genes, as assessed by the pooled CRISPRko screen (p-value and NES were estimated by GSEA 1-tailed test, with 1,000 permutations).

We then reasoned that if MAPK signaling is an important differentiator in all PDLs (with M^+^ states representing the dominant fraction in cell lines), MAPK MRs should be enriched in essential genes that are common across all six PDA cell lines. To test this hypothesis, we generated a CRISPR-based essentiality signature by integrating the signatures of both the CRISPRko and CRISPRi screens across the six PDA cell lines, using Stouffer’s method. As expected, MR generated by VIPER analysis of M^+^ vs. M^-^ cells, integrated across each PDL, were highly enriched in essential genes (*p* ≤ 4.9×10^−5^) (**Figure 5D-E**).

Finally, we used VIPER to identify MRs associated with PDA tumorigenesis by metaVIPER analysis of malignant PDA epithelium (from the CUMC-E dataset), independent of subtype, versus the average of normal tissue samples in GTEx (Consortium, 2013). As expected, PDA tumorigenesis MRs were also significantly enriched in essential genes, as assessed by CRISPRko, integrated across all six PDA cell lines (p = 5.5×10^-3^, by one-tailed GSEA 1000 permutations) (**Figure 5 F-G**). Indeed, GSEA analysis confirmed significant overlap between MAPK and tumorigenesis MRs (p=7.8 X10^-4^, by one-tailed GSEA test with 1000 permutations), thus yielding a subset of MAPK MRs that represent experimentally validated drivers of PDA tumorigenesis (**Figure S4F,** Table S23).

### MR proteins represent mechanistic cell state determinants

While we observed rapid interchange between the M^+^ and M^−^ states in cultured PDA cell lines, the predominance of a single PDL (typically GLS or MOS) in most PDA cell lines limits our ability to assess spontaneous cross-PDL plasticity via barcode-based lineage tracing. Instead, we assessed whether ectopic expression of Lineage-specific MRs could transdifferentiate PDA cell lines from the more aggressive Morphogenic state into the less aggressive Lineage state, thus supporting their mechanistic role in PDL specification. This was assessed by lentiviral-mediated transduction of cDNAs encoding for the 8 most significant Lineage MRs—also among the most activated in the single cell-based GLS state (**Figure S5A**)—in the Morphogenic KP4 cell line using the tetracycline inducible M2rtTA system (Hockemeyer et al., 2008). On an individual basis, ectopic expression of each Lineage MR was effective in increasing the activity of the other Lineage MRs (**Figure 6A**). Yet, ectopic expression of OVOL2 was also uniquely effective in repressing the activity of the top Morphogenic MRs, thus producing virtually complete transdifferentiation of KP4 cells from a Morphogenic to a Lineage state (*p* = 4.3×10^-15^, by 2-tailed aREA test) **(Figure 6A, S5B-C)**.

**Figure 6:**
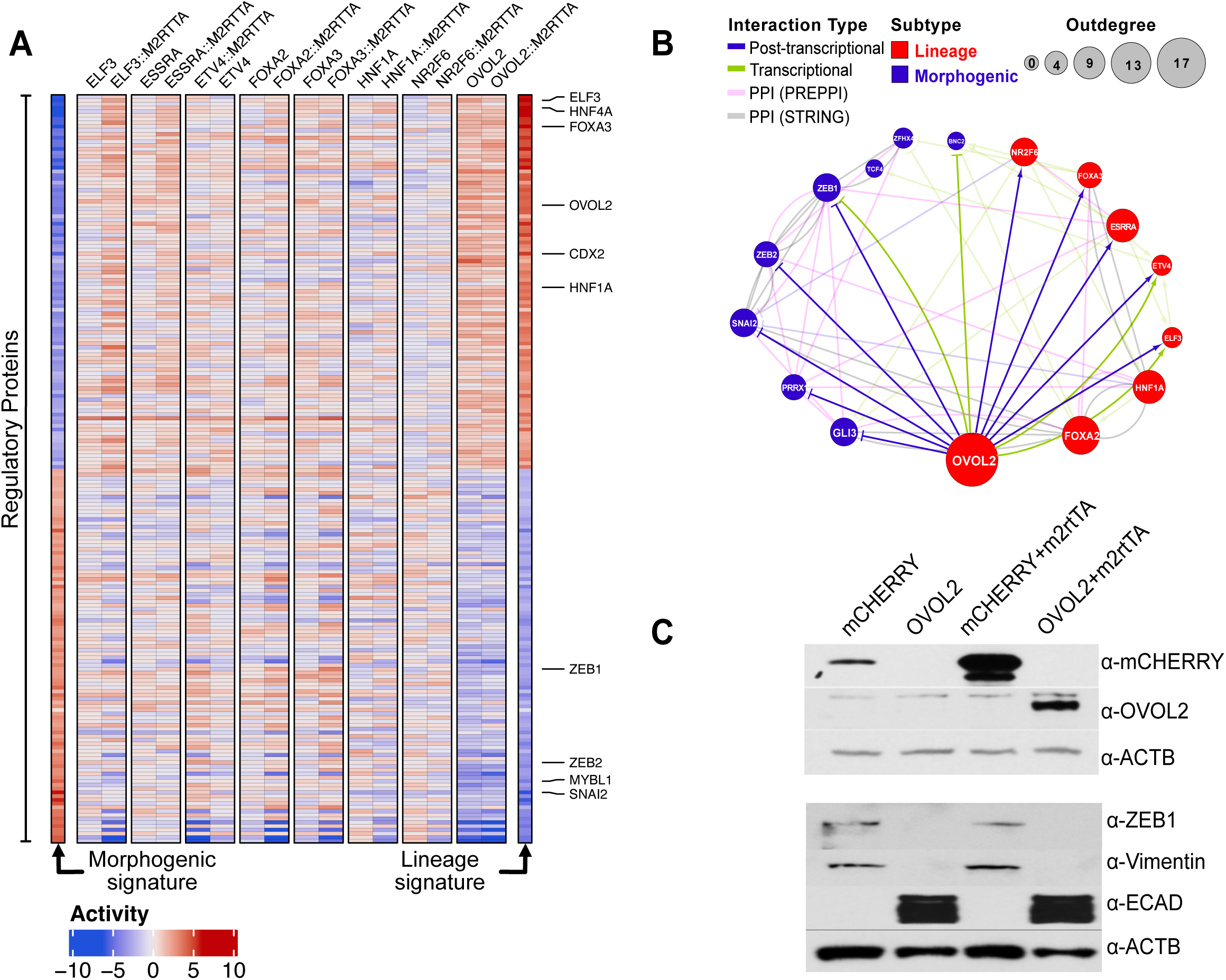
Lineage reprogramming of Morphogenic tumors. **A)** Heatmap showing activity of top Lineage and Morphogenic MRs in the KP4 morphogenic cell line following ectopic expression (+/-M2rtTA) of the top 8 Lineage MRs (n=8). As a reference, to better assess reprogramming, the first and last columns show activity of top Lineage and Morphogenic MRs in KP4 cells (Morphogenic, left) and averaged over HPAFII and PATU cells (Lineage, right). **B)** Mechanistic regulatory network showing transcriptional and post-transcriptional regulation of the 8 top Morphogenic and 8 top Lineage MRs by the latter, as assessed based on their differential expression and differential protein activity analysis following ectopic expression of each MR, respectively. Additional Protein-Protein interactions are reported from the STRING (81) and PrePPI (82) databases. **C)** Western blot showing inhibition of mesenchymal markers (ZEB1 and Vimentin) and expression of epithelial markers (E-cadherin) following ectopic OVOL2 expression (+/-transcriptional activator M2rtTA) in KP4 cells, compared to negative controls (mCherry).

Analysis of RNA-seq profiles following ectopic expression of Lineage MRs in KP4 cells supported reconstructing both transcriptional and post-translational MR → MR interactions. These were assessed by analyzing the differential expression and differential VIPER-measured activity of each MR following ectopic expression of every other MR (see methods). The analysis revealed a complex on/off modular structure where the top 8 Lineage MRs positively regulate each other and repress the top 8 Morphogenic MRs (**Figure 6B**). This modular structure was rich in autoregulatory interactions (loops) contributing to the stability of the two states it regulates, thus providing a mechanistic rationale for the role of these MRs in homeostatic PDL state control. Among the Lineage MRs, OVOL2 emerged at the top of the regulatory hierarchy, due to its ability to directly activate or repress the expression or activity of the vast majority of other Lineage and Morphogenic MRs, respectively, explaining its singular ability to individually control transdifferentiation. Western blotting confirmed OVOL2 cDNA-mediated inactivation of its established transcriptional targets such as ZEB1 and Vimentin (Kang et al., 2018; Zeisberg and Neilson, 2009)—known effectors of EMT, a hallmark of the Morphogenic subtype—and upregulation of established epithelial markers, such as E-cadherin, which is undetectable in KP4 cells (**Figure 6C**). The ability of OVOL2 to transdifferentiate cells to the Lineage state was replicated in in two additional Morphogenic cell lines (PANC1 and PK45H) (**Figure S5D**).

To further dissect the combinatorial logic that drives this Lineage MR module, we performed ectopic cDNA expression of the same top 8 Lineage MRs in individual KP4 cells at MOI = 1, followed by single-cell RNASeq profiling (Son et al., 2021). At this MOI, most cells received one or more cDNAs with co-expression of 11 MR pairs and 14 higher-order MR combinations in ≥ 30 cells (a threshold suitable for transdifferentiation assessment). Consistent with the initial experiment, ectopic OVOL2 expression strongly transdifferentiated cells to the Lineage state. However, the OVOL2/HNF1A pair significantly improved transdifferentiation efficiency by an additional 13% (i.e., 32% to 45.1%) confirming their functional interaction. More importantly, several individual MRs and MR combinations that did not include OVOL2 were also quite effective in inducing Morphogenic → Lineage transdifferentiation in combination but not individually, albeit in a smaller cellular fraction (**Figure 7A-B**). For instance, the FOXA2/HNF1A pair was the third most efficient Morphogenic → Lineage MR combination, even though neither FOXA2 nor HNF1A had a significant effect individually (**Figure 7A**). These results demonstrate the ability of VIPER-inferred Lineage MRs to mechanistically induce the Lineage state, via their transcriptional targets, including as a result of synergistic functional interactions.

**Figure 7:**
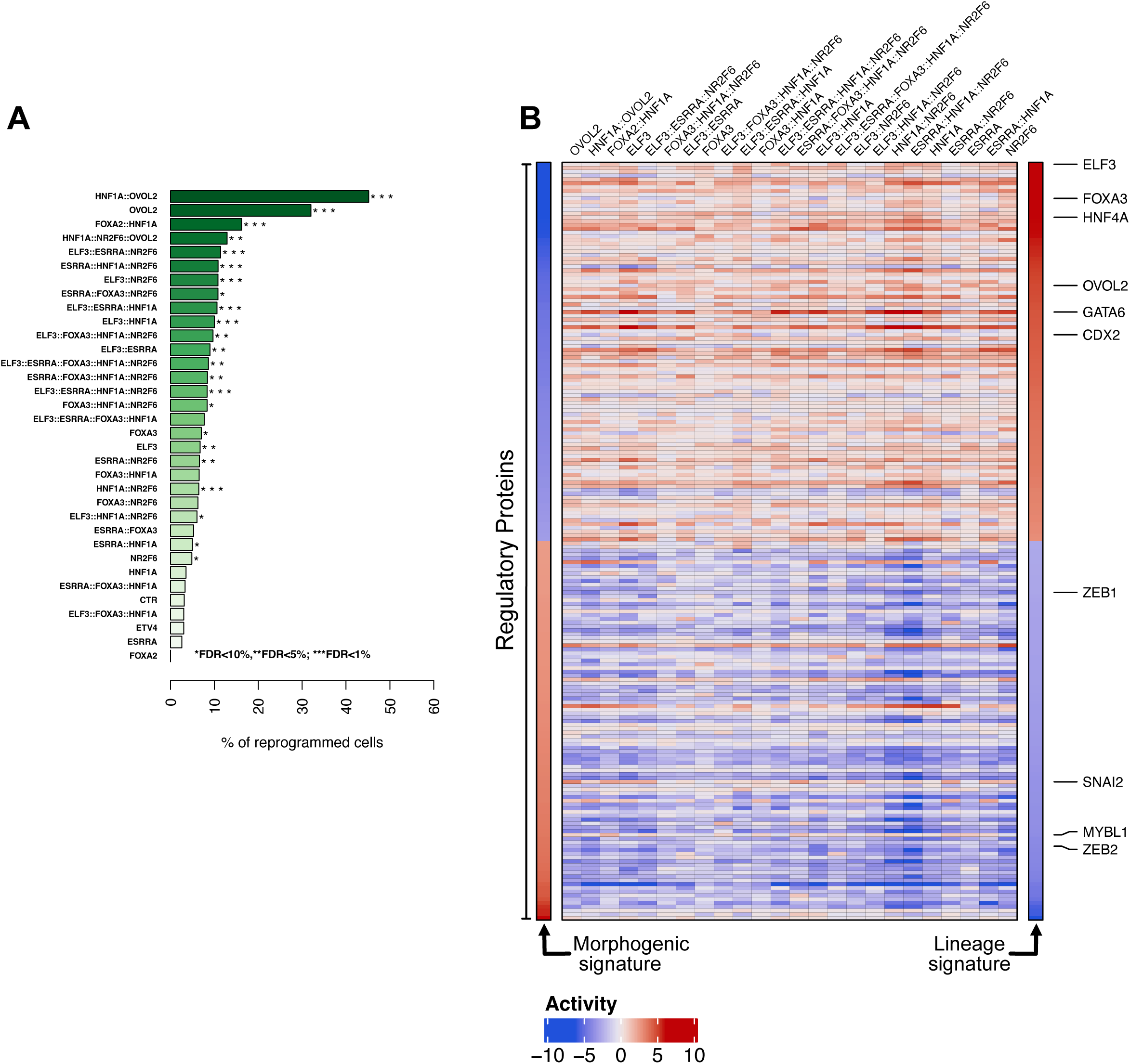
Lineage reprogramming of Morphogenic tumors at single-cell level. **A)** Bar plot showing the fraction of reprogrammed cells following ectopic expression of the top 8 GLS MRs and their combinations in single KP4 cells. P-values were assessed by Fisher Exact Test of the number of statistically significant Lineage cells (FDR<0.05) in KP4 cells, following ectopic expression of individual Lineage MRs and their combination (MR set) vs. negative controls expressing mCHERRY (+/-M2rtTA). **B)** Heatmap showing the average activity profile of Lineage and Morphogenic RPs in the fraction of cells assessed as statistically significantly reprogramed following ectopic expression of each MR set, integrated by the Stouffer ‘s method. Columns represent Lineage and Morphogenic regulators with some notable MRs highlighted.

## Discussion

The parsing of molecular states from bulk PDA expression data has proven extremely challenging due to the wildly heterogeneous composition of these tumors as well as significant variation in patient composition, enrollment criteria, treatment status, and technical variables of available public cohorts. Nevertheless, some consistent top-level messages have emerged, such as the distinction of well-versus poorly differentiated tumors, leading to a growing consideration of subtypes in clinical studies and practice. The assessment of molecular states at the cellular level stands to mitigate issues of intratumoral heterogeneity while revealing the potential contributions of coexisting cellular states to tumor progression and therapeutic resistance. However, two challenges loom before single cell states can be effectively leveraged for therapeutic purposes. First, current techniques for single cell expression analysis suffer from “gene dropout” as low read depth per cell precludes detection of most genes in individual cells. Second, the identification of distinct expression states yields little information regarding the causal drivers that mechanistically implement them. In the current work, we overcome both these challenges using single cell regulatory network analysis, yielding a novel cellular taxonomy, experimental evidence of the mechanisms that implement the respective states, and an understanding of their genetic dependencies for cellular survival.

Enzyme activity assays require assay to quantify products and reactants. For the specific case of enzymes that regulate the abundance of RNA transcripts (e.g., transcription factors, cofactors, chromatin modifiers, and other regulatory proteins), RNA sequencing serves as an ideal assay for activity measurements—*provided* the specific positive and negative target genes of a regulatory protein are known. We rely on ARACNe, an extremely well-validated algorithm based on principles of information theory, to accurately reconstruct the sets of target genes for every regulatory protein in the genome in the specific context of PDA. To avoid the potential biases introduced by reliance on a single dataset, we generated distinct PDA networks from four expression cohorts representing over 500 tumors, including a novel epithelium-enriched cohort from laser capture microdissected samples. Preditions by these networks are then integrated by the metaVIPER algorithm to transform individual expression profiles into regulatory protein activity profiles, yielding powerful benefits such as dimensionality reduction (∼1800 regulatory proteins vs. >20,000 detectable genes), variance stabilization due to the integration of hundreds of gene expression values into one regulatory protein activity value, and virtual elimination of the gene dropout effect. Indeed, a key effect of VIPER analyses is that protein activity can be measured for every regulatory protein on a cell-by-cell basis, *even if the gene encoding for the protein cannot be detected in scRNAseq profile,* making regulatory network analysis ideally suited to single cell datasets. Most critically, the identification of differentially active regulatory proteins explicitly identifies candidate mechanistic drivers of phenotypes, as confirmed by MR-mediated transdifferentiation and CRISPR-ko/CRISPRi studies.

We used this approach to study the malignant epithelial cells of human PDA, leading to identification of six states that were consistently reproduced across multiple single cell expression cohorts. These consisted of three pairs of states that were distinguished by the activities of groups of developmental transcription factors associated with gastrointestinal fate, with acinar-to-ductal metaplasia, or with EMT and morphogen pathway signaling. At the bulk tissue level, the relative abundance of two of these developmental subtypes (GLS and MOS) effectively captured the biology of differentiation state, with the less differentiated Morphogenic tumors exhibiting worse clinical prognosis across multiple independent datasets. Indeed, the conservation of these subtypes across multiple independent datasets with divergent patient and technical properties is a key advantage of this approach.

Critically, the regulatory proteins identified by our work constitute a set of testable predictions concerning the role of each candidate MR in maintaining the identity and viability of their respective cell state. Our experimental validation of these predictions, using ectopic expression assays, represents a critical demonstration of the genetic determinants of cellular plasticity in PDA. Moreover, functional screening confirmed that GLS and MOS MRs are often necessary for maintaining the viability of their respective cell states. These state-specific genetic dependencies provide a roadmap for the future therapeutic targeting of PDA cellular heterogeneity.

Within each pair of developmental states, we identified two substates that were distinguished by their relative level of RAS/MAPK activity. This finding, which was not apparent through the analysis of simple gene expression, is striking considering the nearly universal presence of activating KRAS mutations in human PDA, which might lead to the expectation that all malignant PDA cells have high MAPK signaling. Yet this is clearly not the case as the large majority of malignant epithelial cells in human PDA express low or undetectable levels of the RAF/MEK activity biomarker phospho-ERK by immunohistochemistry. It is well established that mutant KRAS is still ligand-dependent on signaling from upstream receptor tyrosine kinases such as EGFR (Boguski and McCormick, 1993). Local exposure to growth factors, nutrients, and oxygen likely limit the growth and proliferation of malignant PDA cell *in vivo*, leading to far slower proliferative rates compared to replete *in vitro* culture conditions where PDA cells typically divide daily. We hypothesize that the M^+^ and M^-^ states reflect this proliferative heterogeneity. Consistent with this, the large majority of malignant PDA cells are found in the M^-^ state in human tumors, whereas the M^+^ state dominates PDA cells in culture. Treatment with either RAF/MEK/ERK inhibitors or the withdrawal of growth factors (serum free medium), both of which induce cell cycle arrest, effectively shifts PDA cells from the M^+^ to M^-^ state even in the presence of ample nutrients. Nevertheless, the M^-^ state should not be confused with long-term quiescence as our barcode-based single cell fate mapping experiment demonstrates rapid switching between these two states normal growth conditions. Rather, we infer that RAS/MAPK may rapidly toggles during proliferative cycles perhaps in accordance with the metabolic needs of the cell in each cell cycle phase.

The divergence in MAPK state between malignant PDA cells in patients and their counterparts *in vitro* has profound implications for the development of effective therapies. The vast majority of drug screens are performed, at least initially, in cultured cells, potentially biasing for the identification of drugs that selectively target only a small fraction of the malignant cells present *in vivo*. Future screens performed in cells pushed into the M^-^ state may prove more fruitful. Additionally, drugs that selectively impact MRs of just one or two developmental subtypes may produce rapid, non-genetic adaptation to the remaining state. The consideration of regulatory protein activity states may therefore facilitate the development of multi-drug regimens that collectively impact the large majority of malignant cells in PDA.

## ACKNOWLEDGMENTS

This work was supported by a Clinical Translational Program Grant from the Lustgarten Foundation for Pancreatic Cancer Research (KPO) and a Precision Medicine Pilot Award, Irving Institute for Clinical and Translational Research (KPO), NCI Research Centers for Cancer Systems Biology Consortium (U54 CA209997 to AC and KPO), NIH Shared Instrumentation Grants (S10 OD012351 and S1 0OD021764 both to AC), and NCI Cancer Center Support Grant (P30 CA013696) for the Herbert Irving Comprehensive Cancer Center (including the High-Throughput Sequencing, Single-Cell Analysis, OPTIC, Molecular Pathology, and Database Shared Resources). Single-cell studies were supported by the NCI OIA Award (R35 CA197745 to AC). Funding was also provided by The Pancreas Center at Columbia/NY Presbyterian Hospital (KPO), and by the Sigrid Juselius Foundation (MT). We acknowledge support from the Swedish National Genomics Infrastructure, SNIC (project SNIC 2017-7-265), and the Uppsala Multidisciplinary Center for Advanced Computational Science (UPPMAX). We thank Giovanni Valenti for his comments on single-cell subtypes.

## DECLARATION OF INTERESTS

A.C. is founder, equity holder, and consultant of DarwinHealth Inc., a company that has licensed some of the algorithms used in this manuscript from Columbia University. P.L. is sr.Director of Single-Cell Systems Pharmacology at DarwinHealth Inc.. P.L. worked on this project while he was Associate Research Scientist at Columbia University and completed it after he joined DarwinHealth Inc. M.J.A. is CSO and equity holder of DarwinHealth Inc. Columbia University is also an equity holder in DarwinHealth Inc. US patent numbers 10,777,299 and 10,790,040 have been awarded related to this work, assigned to Columbia University.

## AUTHOR’S CONTRIBUTIONS

A.C., K.P.O. and P.L. conceived the project. P.L. started the project, designed, performed and oversaw the computational analyses. M.T. designed, performed and oversaw the experimental analyses. A.C.G., H.C.M, E.E., B.S., J.W., J.K., X.T, E.C.F, K.W., U.N.W., S.G. and F.N. contributed to experimental execution, data acquisition and data generation. L.T., M.J.A., S.T., A.W. and A.G. contributed to the computational analyses. A.I. reviewed histopathology for PDA samples in the CUMC cohort. G.A.M. and W.W. aided in the curation of human outcomes data. A.C., K.P.O. and P.L. wrote the manuscript with feedback from C.H.M, M.T. and M.J.A.. K.P.O., D.A.T. and A.C. supervised the study.

---

**Figure S1:**
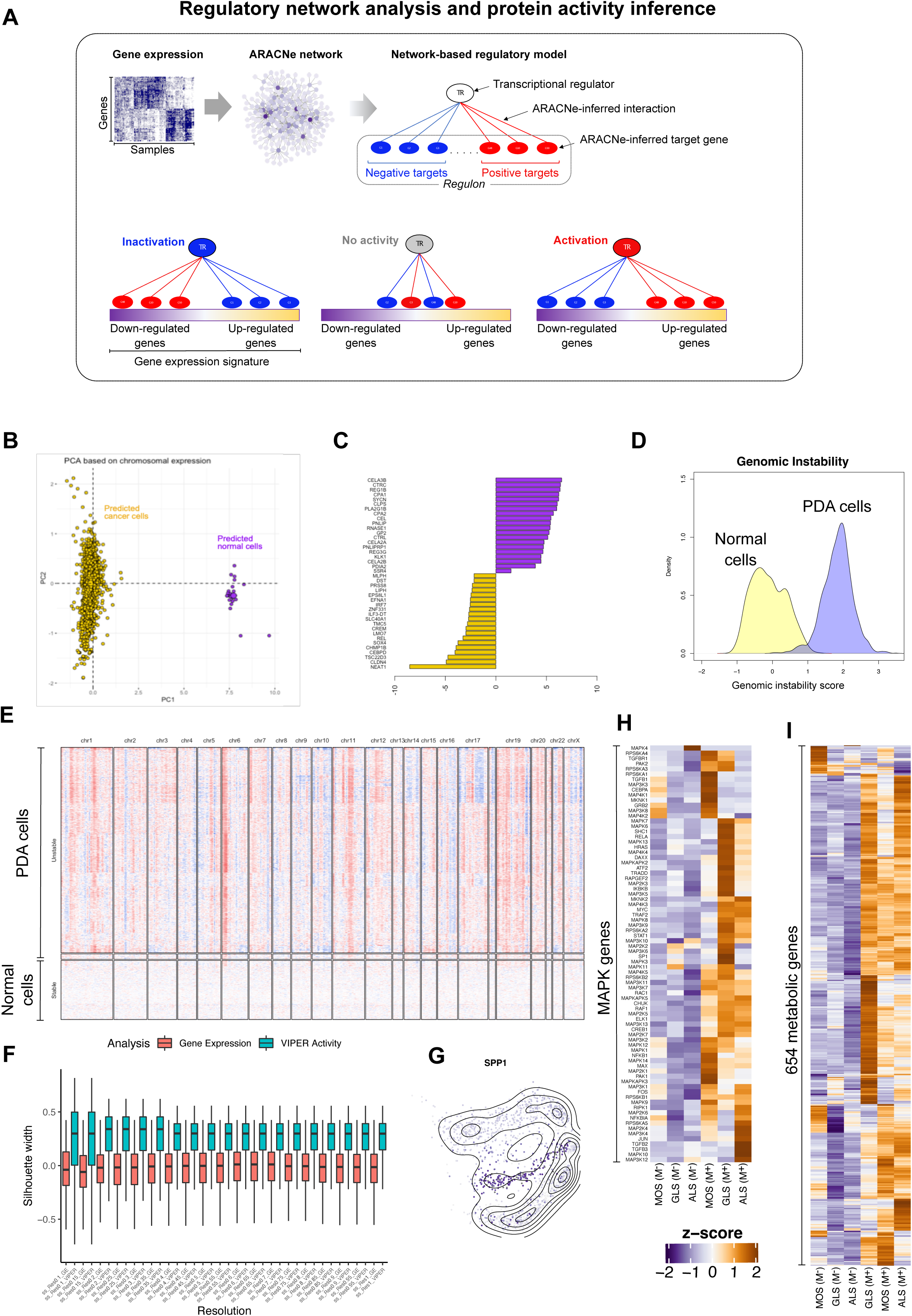
**A)** Conceptual workflow illustrating VIPER-based inference of protein activity. **B)** PCA of single PDA cells based on chromosomal gene expression analysis effectively distinguishes aberrant ploidy tumour cells (yellow) vs. normal-ploidy cells (purple). **C)** Differentially expressed genes between non transfromed (normal) and transformed (cancer) single cells, as predicted by PCA of chromosomal expression. **D)** Density plot showing genetic instability score between predicted PDA cancer cells by PCA on chromosomal expression compared to non transformed pancreatic cells (see methods)**. E)** Heatmap showing CNVs and genomic instability score (G.I. score), as computed from the gene expression profiles of PDA cells compared to non transformed pancreatic cells**. F)** Box plots showing the distribution of silhoutte scores of individual cells clustered by their VIPER-inferred protein actvity profiles and by gene expression at different resolution values of Louvain algorithm. **G)** Plot showing differential gene expression of Spp1 in the first two principal components computed by PCA on protein activity profiles. **H)** Heatmap showing the differential expression of genes associated to MAPK pathways in Biocarta database across the cell states. Differential gene expression is expressed as z-score on the pseudobulk profiles computed by averaging the normalized gene expression profiles (log_2_(CPM+1)) of individual cells within the same cell state (i.e.cluster). **I)** Heatmap showing the differential expression of genes associated to Metabolic pathways in Reactome database across the cell states.

**Figure S2:**
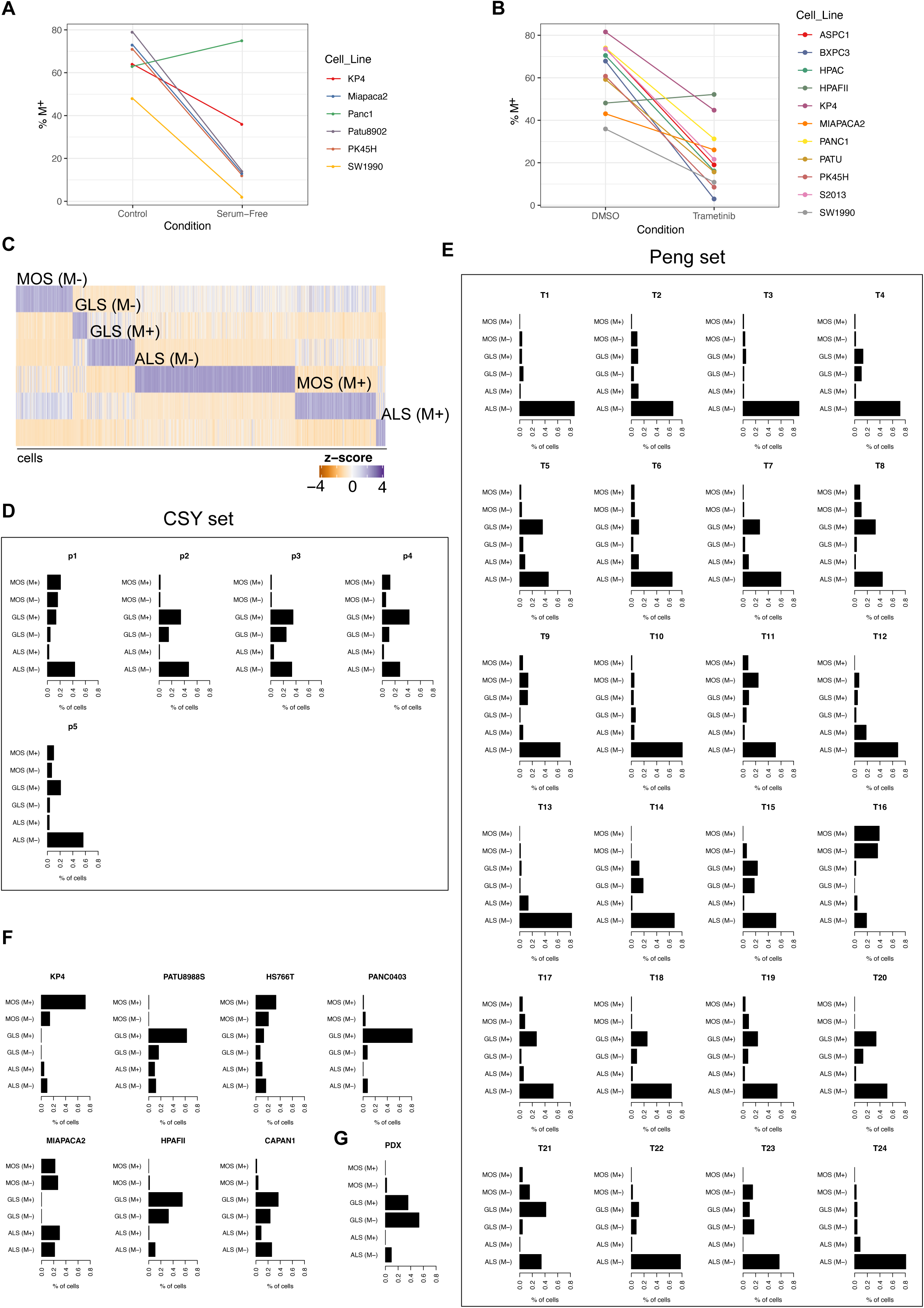
**A)** Plot showing the percent of M^+^ and M^-^ cells in six PDA cell lines after replacing reccomended media (control) with serum-free media. Serum-free media was replaced after 4 hours. Cells were profiled by scRNASeq, transformed to protein activty profiles, and classified as M^+^ or M^-^ based on the enrichment of the experimentally inferred MAPK protein actvity signature. **A)** Plot showing the percent of M^+^ and M^-^ in ten PDA cell lines after treatment with Trametinib with respect the DMSO. Cells were profiled by scRNASeq, transformed to protein activty profiles, and classified as M^+^ or M^-^ based on the enrichment of the experimentally inferred MAPK protein actvity signature **C)** Heatmap showing the differental enrichment of cell state specific protein activity signature (25 most activated and 25 most inactivated) in a PDA patient from CSY set. Enrichment analysis was performd using the aREA algorithm (2-tailed aREA test). **D)** Barplots showing the classification of PDA cells in each patient of the CSY set (single-cell set), based on the enrichment of cell state specific protein activity signature. **E)** Barplots showing the classification of PDA cells in each patient of the Peng set, based on the enrichment of cell state specific protein activity signature. **F)** Barplots showing the classification of PDA cells in seven PDA cell lines, based on the enrichment of cell state specific protein activity signature. **G)** Barplot showing the classification of PDA cells derived from a PDX model, based on the enrichment of cell state specific protein activity signature.

**Figure S3:**
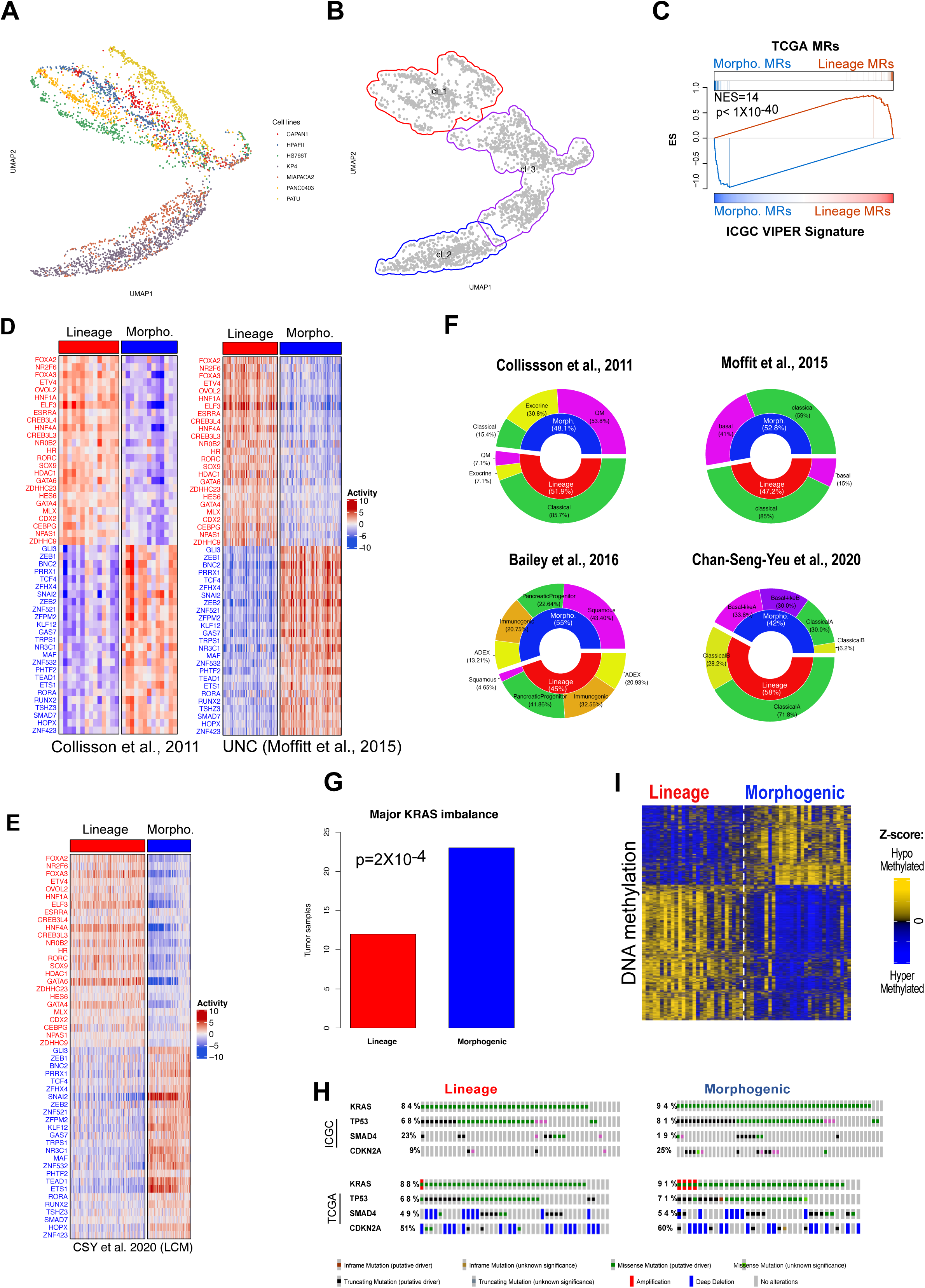
**A)** Reference map (UMAP) based on the protein activity profiles of PDA cells from 7 PDA cell lines. Each dot represents a cell and colors represent different PDA cell lines**. B)** Reference map (UMAP) showing three clusters identified by unsupervised cluster analysis. **C)** GSEA plot showing the enrichment of TCGA MRs (50 most overactivated and 50 most inactivated proteins) in the ICGC protein activity signature (p-value and NES were estimated by GSEA 2-tailed test, with 1,000 permutations). **D)** Heatmaps showing the activity of 25 most activated Lineage and Morphogenic MRs—as obtained by Stouffer integration of ICGC and TCGA sample analysis in the UNC and Collisson cohorts. Only samples with a silhouette scores >0.25 are shown in the heatmaps, which correspond to 102/125 samples (82%) in the UNC cohort and 23/27 samples (85%) in the cohort of Collisson et al.,2011. **E)** Heatmap showing the conservation of 25 most activated Lineage and Morphogenic MRs in a large cohort of 197 laser capture microdissecetd samples from CSY set. **F)** PieDonut charts showing overlap between protein activity-based Lineage and Morphogenic classification and previously published classification schemes. **G)** Barplot showing the enrichment of KRAS inmabalance in Morphogenic patients, in the CSY set. **H)** Oncoprint plot showing genetic alterations in TCGA Lineage and Morphogenic samples from cBioportal (Cerami et al., 2012). **I)** Heatmap showing differentially methylated sites in Lineage vs. Morphogenic TCGA samples (n=58).

**Figure S4:**
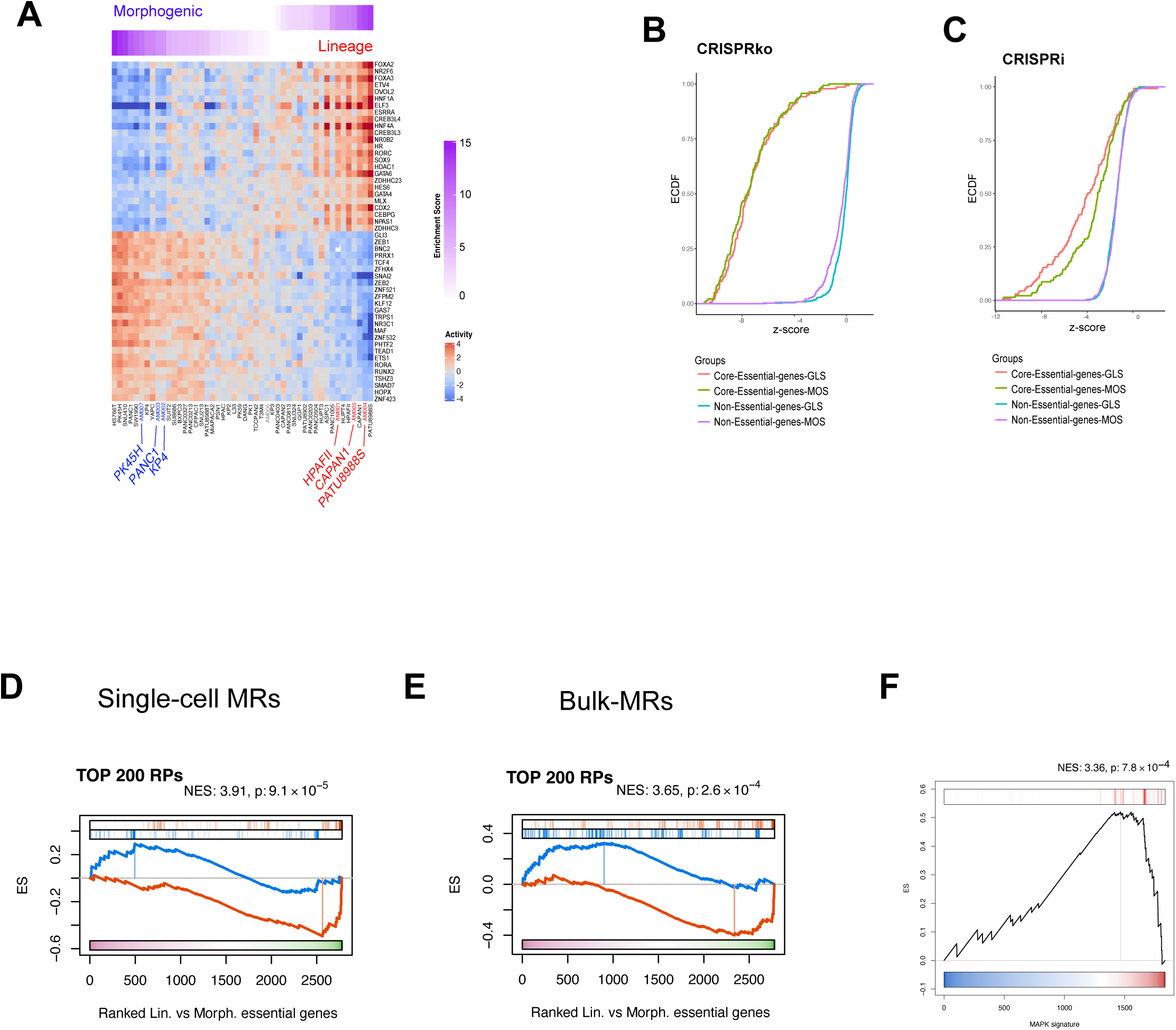
**A)** Heatmap showing the enrichment of the 50 most differentially activated proteins (25 most activated and 25 most inactivated), as inferred by VIPER analysis of Lineage vs. Morphogenic subtype samples, integrated across TCGA and ICGC cohorts, in PDA cell lines from the Cancer Cell Line Encyclopedia (CCLE) (Barretina et al., 2012). The three cell lines labeled in red and blue were selected as Lineage- and Morphogenic specific models, respectively, for a pooled CRISPR/Cas9 screen to validate predicted MR proteins. They were re-sequenced to ensure fidelity compared to their CCLE profiles. Specifically, re-sequenced Lineage cell lines HPAFII, CAPAN1, and PATU8988S were labeled as AM01, AM005, and AM004, respectively, while re-sequenced Morphogenic cell lines PK45H, PANC1, and KP4 were labeled as AM007, AM003, and AM002, respectively. The re-sequenced PANC0403 cell lines (AM006) was classified as neither Lineage nor Morphogenic and was thus selected as control for the cell line selection. **B-C)** For quality control purposes, the ECDF plots show the z-score distribution for established core-essential genes (positive controls) vs. non core-essential genes (negative controls) assessed in the pooled CRISPRko and CRISPRi screens.This shows that the pooled screens were highly effective in identifying core-essential genes, thus supporting the quality of the results. **D)** GSEA plots showing the enrichment of the top 200 differentialy activated proteins (100 most differentially activated and 100 most differentially inactivated) inferred from single cells by comparing GLS vs MOS states, in the Lineage vs. Morphogenic essential genes. P-value and NES were estimated by two-tailed GSEA test with 1000 permutations. **E)** GSEA plots showing the enrichment of the top 200 differentialy activated proteins (100 most differentially activated and 100 most differentially inactivated) between Lineage and Morphogenic subtypes inferred from bulk anlysis (ICGC-TCGA signature), in the Lineage vs. Morphogenic essential genes. P-value and NES were estimated by two-tailed GSEA test with 1000 permutations. **F)** GSEA plot showing the enrichment of the 50 most overactivated RPs of the PDA tumorigenic signature (inferred by VIPER analyss by comparing CUMC-E vs GTEx) in the MAPK signature inferred by VIPER analysis from single cells of PDA patients. NES and p-value were estimated by one-tailed GSEA test with 1000 permutations.

**Figure S5:**
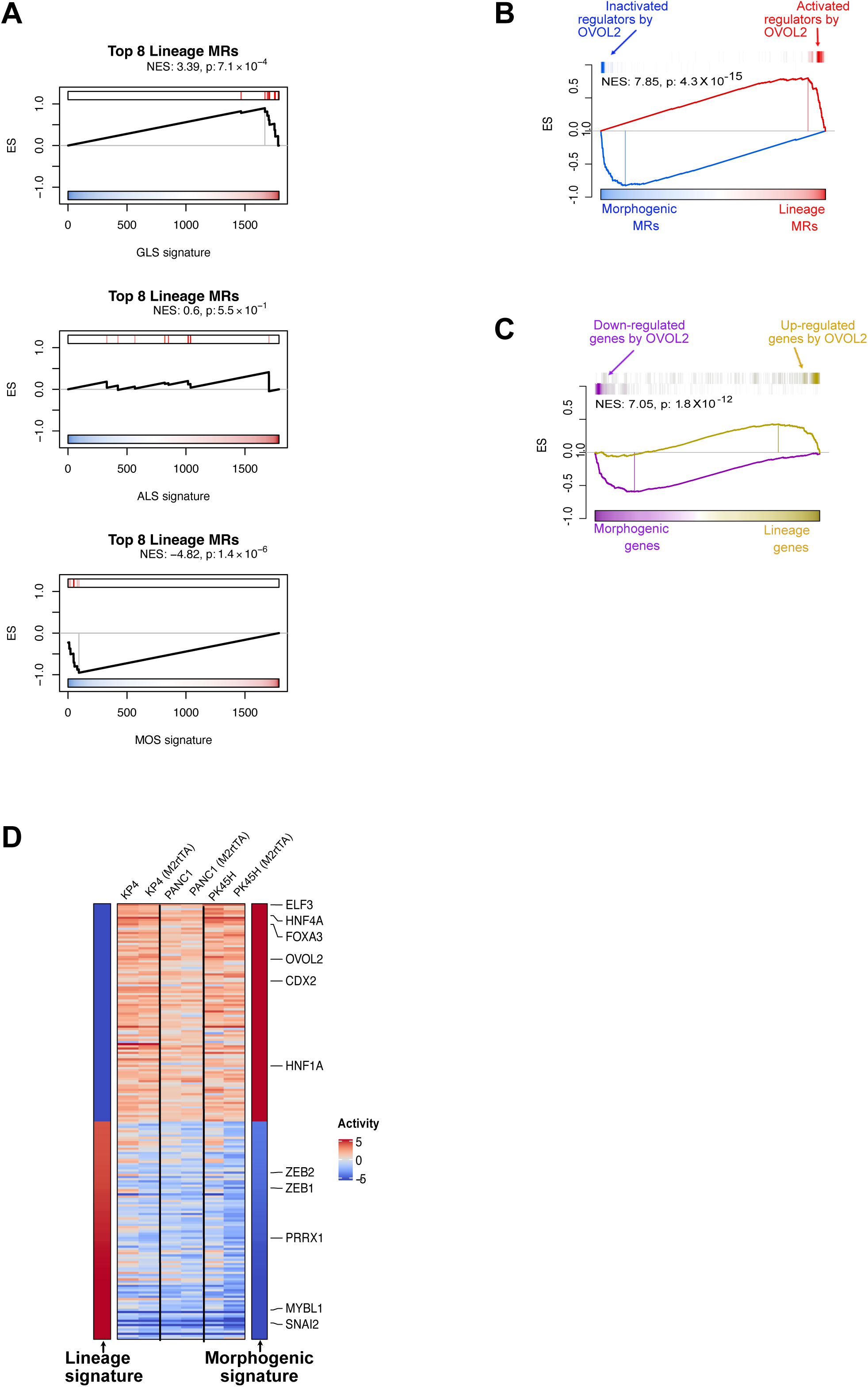
**A)** GSEA plots shwoing the enrichment of the top eight most over-activated Lineage RPs in the single-cell PDL signatures computed by comparing each PDL against the other two. These plots show the conservation of the top eight Lineage RPs in the GLS signature. NES and p-value were estimated by two-tailed GSEA test with 1000 permutations. **B)** GSEA plot showing the enrichment of the top 50 Lineage (red) and top 50 Morphogenic (blue) MRs in proteins differentially activated following ectopic OVOL2 expression in KP4 cells. P-value and NES were assessed by two-tailed GSEA test with 1000 permutations. **C)** GSEA plot showing the enrichment of 200 most over-(yellow) and under-expressed (purple) genes following ectopic OVOL2 expression in KP4 cells in genes differentially expressed in Lineage vs. Morphogenic cell lines (p-value and NES assessed by two-tail GSEA analysis with 1000 permutations). **D)** Heatmap showing reproducibility of OVOL2-mediated Morphogenic → Lineage cell state reprogramming (+/-M2rtTA), across three distinct Morphogenic cell lines (KP4, PANC1 and PK45H).

## METHODS

### ARACNe networks

PDA networks were generated from CUMC, ICGC and UNC cohorts using the ARACNe-AP algorithm (Lachmann et al., 2016) with 100 bootstrap iterations and a mutual information (MI) *p*-value threshold of 10^-8^, corrected for multiple hypothesis testing, as originally described (Basso et al., 2005; Margolin et al., 2006). TCGA networks, including the PDA TCGA network, were downloaded from the aracne.networks package (Federico M. Giorgi). ARACNe networks included transcriptional targets (regulons) of a set of Regulatory Proteins comprising TFs, co-TFs, and chromatin regulators was used for this study (Table S1).

### Single-cell analysis of the Elyada PDA set

The single-cell UMI-count matrix from Elyada et al.(Elyada et al., 2019) (Elyada set) was filtered to remove cells with <1,000 UMI-counts and genes with zero counts across all cells. UMI counts were normalized to counts per million (CPM). Epithelial cells were computationally selected from each sample using a GSEA clustering procedure based on enrichment of cell type specific markers, including those for epithelial, endothelial, immune, fibroblasts and pericytes cells (see supplementary methods) as also discussed in (Elyada et al., 2019). 1886 cells from six patient-derived samples were identified as epithelial. Of these, 30 cells were further removed as putative non cancer cells as predicted by genomic instability (Laise and Alvarez, 2022) and aneuploidy analyses(Yuan et al., 2018). For genomic instability analysis single cells derived from normal pancreas were used as a reference (Han et al., 2020). 1856 cells were identified as putative cancer epithelial cells. A representative subset of 500 cells—selected at random from the 1,856 available PDA epithelial cells—was used to build a single-cell ARACNe network (scNET). CPM normalized counts were used with 100 bootstrap iterations and a Bonferroni corrected statistical significance threshold, p ≤ 10^-8^. ARACNe inferred adequately sized regulons (i.e., >50 targets) for 506 of 1,835 regulatory proteins. For the remaining regulatory proteins (n = 1,329), activity was inferred by integrating four PDA networks (CUMC-net, TCGA-net, ICGC-net and UNC-net) via the metaVIPER algorithm. All ARACNe regulons were pruned to the 50 most-statistically significant targets before metaVIPER analysis, to avoid bias associated with different regulon sizes being used in the analysis. To compute a differential gene expression signature for each individual single-cell (for metaVIPER analysis) we used the “mad” method—similar to a robust z-score (Malo et al., 2006)—as implemented in the VIPER package (Alvarez et al., 2016), on the rank transformed single cell RNASeq (scRNASeq) profiles. Cluster analysis was performed on the metaVIPER inferred protein activity profiles using the Louvain algorithm as implemented in the Seurat package (Stuart et al., 2019) in the viper space computed using the viperSimilarity function of the VIPER package (Alvarez et al., 2016) The viperSimilarity function computes the distance between each pair of cells by performing the reciprocal enrichment analysis (two-tailed aREA test (Alvarez et al., 2016)) of the protein activity signatures, and generates a distance matrix based on the similarity of the protein activity signatures. This distance matrix was used as input to construct a Shared Nearest Neighbor (SNN) graph using the “FindNeighbors” function of the Seurat package with the “k.param”=50. The optimal number of clusters was estimated by optimizing the resolution parameter (from 0.1 to 1 with intervals 0.05) of the “FindNeighbors” function with silhouette analysis (Rousseeuw, 1987) This analysis estimated 0.2 as optimal resolution value, which generated 6 clusters. A differential protein activity analysis was then performed to generate a protein activity signature for each cluster. This was done using the most representative cells of each cluster selected based on the silhouette scores (n=100 cells with the highest silhouette score in each cluster were selected). The differential protein activity analysis was performed by metaVIPER on the differential gene expression signatures computed by comparing the most representative cells of each cluster against the most representative cells of all the other clusters (one vs. all) using the Student T.test on the Log2 (CPM+1) normalized gene expression profiles. Pseudo trajectory analysis was performed on the VIPER-inferred protein activity profiles (computed using the robust z-score as previously described) of all cancer cells using the Monocle algorithm (Trapnell et al., 2014).

### Drug perturbation assays

PLATE-Seq experiment was performed in collaboration with Columbia University’s Genome Center. Panc1 and Aspc1 pancreatic cancer cells were cultured in white 96-well tissue culture-treated plates at optimized density, in 100 μl of Dulbecco’s Modified Eagle Medium (DMEM) media supplemented with 10% fetal bovine serum (FBS) and 1% penicillin/ streptomycin. After 24 h of incubation, the plates were treated with following drugs: RAF inhibitors – Sorafenib, Dabrafenib, RAF709, PLX8394, GDC-0879; MEK inhibitors – Trametinib, Cobimetinib, Binimetinib, Selumetinib, Rafametinib; and ERK inhibitors – SCH772984, Ulixertinib, AZD0364, Ravoxertinib. Each drug was dosed at the concentration at which the cells were 80% viable after 48 h of treatment. After 24 h of treatment, the medium was replaced with 100 ml of FBS supplemented with 10% DMSO and the plates were frozen at −80 °C prior to PLATE-Seq. Detailed protocol for preparation of the automated PLATE-SEQ experiment was described by Bush et al. (Bush et al., 2017). The PLATE-Seq FASTQ files were pseudoaligned to the GRCh38 human transcriptome (mRNA & ncRNA) and gene expression was quantified using kallisto (version 0.44.0), tximport package (Soneson et al., 2015), and biomaRt package (Durinck et al., 2009). The gene expression was quantified as both raw counts (i.e. sequencing fragments per genomic locus) and transcripts per million (i.e. sequencing fragments per genomic locus normalized for transcript/gene length and sample sequencing depth). ShortRead package(Morgan et al., 2009) was used to assess the quality of sequencing data for each sample. The number of total sequenced reads, the number of aligned sequencing reads, and the read alignment proportion for each sample were assessed. Single sample differential gene expression signatures were computed independently for each one of the two cell lines. The z-score method was used to generate differential gene expression signatures of each drug-treated sample with respect to the DMSO-treated samples (reference). Protein activity profiles were computed using the metaVIPER approach as in “Single-cell analysis of the Elyada PDA set”.

### Serum Free media treatment

Six human PDA cell lines (KP4, SW1990, PK45H, Panc1, MiapaCa2 and Patu8902) were seeded in a 6-well plate (Falcon, ref. 353046) using recommended media. 4h later, media was changed or replaced for serum-free media and culture them for 48h. Cells were then processed for single cell RNAseq as described in (Peng et al., 2019) using Total Seq B antibodies and the recommended protocol from Biolegend. Briefly, a digestion buffer that contained trypsin, DNAse and enzymatic cocktail (Miltenyi, Cat. No. 130-095-929) and the gentleMACS Octo Dissociator (Milteny Biotec, Cat. No. 130-095-937) were made for initial tumor disruption using manufacture’s protocol. Cell suspensions were then filtered using a 40μm cell strainer (Falcon, Cat. No. 352340) and red blood cells (RBC) were removed by RBC lysis buffer (Invitrogen, Cat. No. 1966634). Dissociated cells were washed twice with PBS 0.1% BSA buffer with cold centrifuging at 500rpm and 5’. Finally, cells were stained with 0.4% Trypan blue (Invitrogen, Cat. No. T10282) to check the viability and diluted with PBS 0.1% BSA to about 1E106 cells/ml for single cell sequencing. For each single cell run, 6 samples were combined. Single cell data were filtered for low quality cells and normalized as previously described in the “Single-cell analysis of the Elyada PDA set”. To assess the effect of serum free media on the M^+^ and M^-^ PDA cells, we first computed the number of M^+^ and M^-^ cells in the control cells (cells in the recommended media). This was done by applying the metaVIPER approach on the single cell gene expression signatures (z-score transformed gene expression profiles) of the control cells computed using as reference the gene expression profiles of seven PDA cell lines (see single-cell cross cohort analysis). The count data of the seven PDA cell lines were downsampled to make the probability distributions of the number of UMIs per cell comparable across different experiments. The number of M^+^ and M^-^ cells was computed by assessing the enrichment of the top 100 MRs (top 50 most activated and top 50 most inactivated) of the MAPK signature inferred in the perturbation assay. Then, we assessed the effect of the Serum free media by comparing single cells of each cell line between the two experimental conditions (i.e., serum free media vs control cells), using the metaVIPER approach. Cells were classified as M+ or M-base on the enrichment of the MAPK MRs as described above.

### Trametinib treatment

For single cell analysis of MAPK pathway inhibition, 11 PDA cell lines were used. Optimal concentrations to inhibit MAPK pathway were specific to each cell line and they were optimized using pERK Western Blot analysis (not shown). Cells were seeded on 6 well plates, and on the next day treated with Trametnib. After 24h of incubation cells were collected for single cell analysis. Cells were then processed for single cell RNAseq as described in (Peng et al., 2019). For each single cell run, 8 samples were combined. Each pre-incubated with a human specific barcode. Vehicle and treated for each cell line were run in the same lane. Viability for each sample was above 80%. Single cell data were filtered for low quality cells and normalized as previously described in the “Single-cell analysis of the Elyada PDA set”. To assess the effect of Trametinib treatment on the M^+^ and M^-^ PDA cells we used the same approach described for the analysis of serum free media treatment, with DMSO treated cells as control cells.

### Single cell cross-cohort analysis

Cross cohort consistency of the protein activity signatures identified in the Elyada set was performed in two independent PDA human cohorts, including the CSY-scSet (Chan-Seng-Yue et al., 2020)(n=5 patients) and Peng set (Peng et al., 2019) (n=24 patients), in 7 PDA cell lines (CAPAN1, HPAFII, HS766T, KP4, MIAPACA2, PANC0403, PATU8988S), and in a PDA PDX model. The counts matrices for all the data sets were filtered for low quality cells as described in the analysis of Elayada set. For the CSY-scSet, the count matrix was made available by the Notta lab. The Peng data were downloaded from the Genome sequencing Archive as reported by the authors. Single cell RNAseq data for the 7 PDA cell lines were generated as following: Cell lines were trypsinized and resuspended in Cell Staining Buffer (Biolegend, Cat. No. 420201) and incubated with proper Total Seq B antibodies using recommended protocol from Biolegend. Cells were washed twice with PBS-0.5%BSA solution and mixed them together aiming at even cell numbers for each cell line. To assess whether the confluence state of cell line would affect its protein activity profiles and, consequently, its classification, the KP4 cell line was profiled at high-confluence states (KP4-HC) and low-confluence (KP4-LC). Single-cell sequencing data were processed using the Cell Ranger pipeline (v.5.0.1) from 10X GENOMICS (https://www.10xgenomics.com/). FASTQ files were aligned using the human genome as a reference (v. GRCh38-2020-A). The combination of Cite-seq-Count (https://github.com/Hoohm/CITE-seq-Count) and the *HTODemux* pipeline of the Seurat package was used to demultiplex the data and assign each single-cell to corresponding cell line of origin, as explained in (Stoeckius et al., 2018).

All the cells that resulted to be positive to more than 1 antibody or negative for all of them were not included in the downstream steps of the analysis. The barcode sequences of the BioLegend antibodies that were used to tag each cell line are listed supplemental methods. PDA tumors from PDX mice were dissociated using the protocol described in (Peng et al., 2019) (see “Serum Free Media Treatment” for more details). For the PDX model, single-cell sequencing data were processed using the Cell Ranger pipeline (v.3) from 10X GENOMIC (https://www.10xgenomics.com/). FASTQ files were aligned on both GRCh38-3.0.0 and gex-mm10-2020-A transcriptomes. Differential mapping analysis was performed for each cell, by comparing the number of reads mapped to the human vs. mouse transcriptome. Cells that had ≥ 25% more reads mapped on human transcriptome than mouse transcriptome were considered as human tumor cells and used for downstream analyses.All the count matrices were filtered for low quality cells, normalized to CPM and analyzed independently. The normalized gene expression matrix of the PDA cell lines was transformed into a gene expression signatures matrix using the z-score transformation and then transformed into protein activity profiles using the metaVIPER approach as described in the analysis of the Elyada set. The count matrix of the PDX model was transformed in a gene expression signature matrix by z-score transformation using the Elayda set as reference and then transformed into protein activity profiles using the metaVIPER approach as described in the analysis of the Elyada set. The reproducibility/conservation the protein activity signatures identified in the Elyada set in the other independent data sets was computed by performing a two-tailed enrichment analysis using the aREA algorithm(Alvarez et al., 2016; Alvarez et al., 2019) of the 50 most differentially activated regulatory proteins (25 most overactivated and 25 most inactivated) in the protein activity profile of each individual cell.

### Single-cell Lineage Tracing

We modified the lentiviral vector also used in our transcription factor overexpression assays (modified Tet-O-FUW EGFP-puro, addgene #30130) by using mCherry instead of EGFP, and cloned a UMI-barcode (UMI-bc), which consists a random 28-mer, 200 bp upstream of the lentiviral 3’-long terminal repeat (LTR) region. This way the UMI-bc site will get polyadenylated and barcode can be specifically amplified in the end of scRNA-seq library preps. In total we cloned a plasmid library which contains approx. 6 million distinct UMI-barcodes. The basic principle for lineage tracing experiments was similar as in (PMID 31974159). Briefly, the UMI-bc containing lentiviral constructs were transduced into KP4 and PATU8988S cell lines (to approx. 15-30 million cells) with MOI <0.1 followed by puromycin selection. After all the non-transduced cells were dead, we seeded 8000 cells / well, followed by incubation which lasted approx. 1-2 population doublings (daughter cells were created for each UMI-BC containing cell). At this point we performed the 1^st^ chromium run (time point 1) so that 50% of the cells were used for this this initial time point run, and the remaining 50% of the cells were cultured further for the 2^nd^ time point. The 2^nd^ time point chromium run was done approx. 10 population doublings later. For both the time points and both the cell lines, the entire pool of UMI-BCs was processed using UMI-tools (https://www.ncbi.nlm.nih.gov/pmc/articles/PMC5340976/) to collapse clusters of UMI-BCs with less than 4 mismatches. Through this procedure, a reference list of all detected UMI-BCs was generated, and it allowed us to identify all the cells associated to one unique UMI-BC.

### Analysis of Single-cell Lineage Tracing data

Single-cell sequencing data of the PATU8988S and the KP4 cell lines were processed using the Cell Ranger pipeline (v.3.0.2) from 10X GENOMICS (https://www.10xgenomics.com/). FASTQ files were aligned using the human genome as a reference (v. GRCh38-2020-A). In order to avoid differences in sequencing depth across cells sequenced in different runs, which could significantly affect gene detection, the UMI counts of PATU8988S, KP4 and the other 7 PDA cell lines were downsampled to make the probability distributions of the number of UMIs per cell comparable across different experiments.

Single-cell gene expression profiles generated from PATU8988S and KP4 cell lines were normalized to CPM and transformed to gene expression signatures using the gene expression centroid of the 7 PDA cell lines as reference (z-score transformation). Gene expression signatures were transformed into protein activity signatures using the metaVIPER approach as described in “Single-cell analysis of the Elyada PDA set”. All the downstream analyses were then performed on the metaVIPER inferred protein activity profiles. A PCA followed K-means clustering, with silhouette analysis for estimating the optimal number of clusters, was performed on the refence cells. The top 30 PCA components were then used to generate a reference map (UMAP). The reference map showed a clear bifurcation separating GLS and MOS cells, while cluster analysis identified 3 centroid-based clusters with two clusters clearly separating GLS (M^+^) and MOS (M^+^) cells and one cluster representing the bifurcation point comprising of M^-^ cells committed toward the MOS (M^+^) or GLS (M^+^) clusters. Then, allowing us to separate KP4 MOS (M^+^) from MOS (M^-^), and PATU8988S (M^+^) from PATU8988S (M^-^). Protein activity profiles of KP4 and PATU8988S cells generated for lineage tracing were mapped in the PCA space of reference cells using the “predict” function of “stats” library of R programming language, and classified as (M^+^) or (M^-^) based on a KNN algorithm trained on the first two principal components (PCA components) of the reference cells using the “train.knn” function of the traineR package (available on https://cran.r-project.org/). The top two components were selected based on the elbow method. The predict function was used to project the KP4 and PATU8988S single cells from lineage tracing experiments in the UMAP space of the reference cells.

### Laser Capture Microdissection data set (CUMC-E)

Freshly frozen tissue samples were obtained from patients who underwent surgical resection at the Pancreas Center at Columbia University Medical Center as previously described (Maurer et al., 2019). Prior to surgery, all patients had given surgical informed consent, which was approved by the institutional review board. Immediately after surgical removal, the specimens were cryopreserved, sectioned and microscopically evaluated by the Columbia University Tumor Bank (IRB AAAB2667). Suitable samples were transferred into OCT medium (Tissue Tek) and snap frozen in a 2-methylbutane dry ice slurry. The tissue blocks were stored at −80°C for later processing. H&E stained sections of frozen PDA samples from the Tumor Bank were initially screened to confirm diagnosis and overall sample RNA quality was assessed by the Pancreas Center supported Next Generation Tumor Banking program using gel electrophoresis, with samples exhibiting high RNA quality utilized for subsequent analyses, including: Laser Capture Microdissection (LCM), RNA sequencing and gene expression quantification. LCM-RNASeq was performed as previously described (Maurer et al., 2019; Maurer and Olive, 2019). Briefly, Cryosections of OCT-embedded tissue blocks were transferred to PEN membrane glass slides and stained with cresyl violet acetate. Adjacent sections were H&E stained for pathology review. Laser capture microdissection was performed on a PALM MicroBeam microscope (Zeiss), collecting at least 1000 cells per compartment. RNA was extracted and libraries prepared using the Ovation RNASeq System V2 kit (NuGEN). Libraries were sequenced to a depth of 30 million, 100bp, single-end reads on an Illumina HiSeq 2000 platform.

### Analysis of publicly available PDA RNASeq data sets and of CUMC-E datset

RNASeq gene counts from CUMC, ICGC (Bailey et al., 2016) and TCGA (https://www.cancer.gov/tcga) cohorts were normalized by variance stabilization transformation (VST), as implemented in DESeq2 package (Love et al., 2014). To avoid excessive stromal contamination as a confounding factor, we selected only samples annotated as “high purity” in the TCGA cohort. Microarray data from Collisson et al.,(Collisson et al., 2011) and Moffitt et al.,(Moffitt et al., 2015) were downloaded as normalized gene expression profiles. A differential gene expression signature for each sample/patient was generated independently for each cohort from the normalized gene expression profiles using the “*scale”* method *(z-score)* implemented in the VIPER package (Alvarez et al., 2016). Differential gene expression signatures of CUMC cohort samples were transformed into protein activity profiles using the VIPER algorithm (Alvarez et al., 2016), leveraging a PDA specific ARACNe regulatory network generated from the epithelial compartment gene expression profiles of CUMC-E samples (CUMC-net). The rationale was to generate a reference data set of protein activity profiles from pure epithelial samples to asses whether the findings obtained by applying the more generalizable metaVIPER approach would be recapitulated in the less generable but biologically relevant CUMC-E cohort. The same network was used to generate the protein activity profiles of a second PDA LCM epithelial gene expression data set profiled in, (Chan-Seng-Yue et al., 2020) and made available as count matrix by Notta lab. Differential gene expression signatures for the other PDA cohorts and PDA cell lines were transformed into protein activity profiles using the metaVIPER approach (Ding et al., 2018) Cluster analysis was performed independently in each cohort by applying the Partitioning Around Medoids algorithm (PAM) as implemented in the cluster package (Martin Maechler, 2019), using a VIPER-based distance metric. Specifically, the VIPER distance between two samples is computed using the reciprocal (i.e., integration of both direct and reverse) enrichment analysis (Kruithof-de Julio et al., 2011) of the Tumor Checkpoint proteins (i.e., 25 most activated and 25 most inactivated) in one sample in proteins differentially activated in the second sample, as implemented by the *viperSimilarity* function in the VIPER package (Alvarez et al., 2016). Use of 50 proteins (defined as Tumor Checkpoint protein) for sample similarity analysis is based on recent results showing that, on average, across all TCGA cohorts, the top 50 most aberrantly differentially activated proteins (candidate Master Regulators) are sufficient to canalize the effect of >90% of somatic mutations, on a sample by sample basis (Paull et al., 2020). Optimal cluster number was then estimated based on the global similarity of all samples in a cluster (cluster membership strength)—as computed based on the conservation of differential protein activities across all samples in the cluster—and evaluated by an Area Under the Curve (AUC) metric (Till, 2001). The optimal number of clusters was also evaluated as by Silhouette analysis (see supplementary methods for details).

### Immunohistochemistry for phospho-ERK

All staining was performed on 4µm paraffin sections of human tissue. For immunohistochemistry, sections were deparaffinized and antigen retrieval was performed in a pressure cooker for 5 min in 1X sodium citrate buffer, pH 6.0 (Abcam). 3% H2O2 was used to block endogenous peroxidases. Slides were then blocked in 2.5% horse serum for 1hr and then incubated in anti-phosphoERK primary antibody (Cell signalling, 1:200) overnight at 4C. The next day, slides were washed in 1X PBS-T and incubated with anti-rabbit secondary antibody for 30 min (Vector Laboratories). Following incubation, slides were washed with 1X PBS-T, developed with ImmPACT DAB peroxidase (Vector Laboratories), and counterstained with hematoxylin.

### Survival analysis

Survival analysis was performed by comparing samples in different protein activity-based clusters, using the Kaplan-Meier method, as implemented in the R “survival” software package (Therneau, 2020). P-values were computed by log-rank test. Kaplan-Meier curves were generated using the “survminer” software package(Alboukadel Kassambara, 2019)

### DNA methylation analysis

450K DNA methylation profiles were downloaded from TCGA using TCGAbiolinks package (Colaprico et al., 2016). Beta values were converted to M-values using the “beta2m” function implemented in the Minfi package (Aryee et al., 2014). Differential methylation analysis between Lineage and Morphogenic samples was performed on M-values using the limma package (Ritchie et al., 2015). All probes with a FDR<0.05 were considered as differentially methylated. A cluster analysis based on differentially methylated sites was performed using PAM algorithm and evaluated by silhouette analysis. Only samples with a positive silhouette score were represented in the DNA methylation heatmap (n=58/76).

### Identification of cell lines representative of Lineage and Morphogenic PDA subtypes

RNASeq count data were downloaded from the Cancer Cell Line Encyclopedia (CCLE) portal (https://portals.broadinstitute.org/ccle) and normalized using the VST method (Love et al., 2014). Differential gene expression signatures for each cell line vs. the average of all cell lines was first computed using the “scale” method in the VIPER package (Alvarez et al., 2016) and then transformed into differential protein activity signatures using metaVIPER (Ding et al., 2018). Cell lines representative of Lineage and Morphogenic subtypes were identified based on the enrichment of Lineage-Morphogenic Tumor Checkpoint proteins in protein differentially active in each cell line, by two-tailed aREA test (Alvarez et al., 2016).

### RNA extractions and re-sequencing of PDA cell lines representative of Lineage and Morphogenic subtypes

Selected PDA cell lines were cultured in 6-well plates such that confluency at 48h-72h post seeding was < 50%. Total RNA was extracted with the RNeasy Plus mini kit (Qiagen) and sequenced on the NovaSeq 6000 (PE 20million reads). Reads were processed using the Kallisto pipeline(Bray et al., 2016), with GRCh38 as reference.

### Protein activity analysis of re-sequenced Lineage and Morphogenic representative cell lines

First, RNASeq count profiles of re-sequenced cell lines were added to the CCLE count matrix; then, the count matrix was VST normalized (Love et al., 2014) and differential gene expression signatures were generated with the “*scale”* method *(z-score*) in the VIPER package; finally, differential gene expression signatures were transformed to differential protein activity profiles using metaVIPER (Ding et al., 2018). Cell line subtype was assessed based on the enrichment of Lineage-Morphogenic Tumor Checkpoint proteins in proteins differentially active in each cell line, by two tailed aREA test (Alvarez et al., 2016).

### CRISPRko and CRISPRi screening

sgRNA containing lentiviruses were transduced into Cas9 expressing PDA cell lines in duplicates (in presence of 8ug/ml polybrene), at an estimated MOI = 0.2 - 0.3. After 24h, the lentivirus containing media was removed, cells were washed with PBS, and puromycin-containing media (2ug/ml) was added to the cells for 48-72h until all control cells (not virus-infected) were dead. Half of the cells were harvested at this time point (day0) and sequenced to assess sgRNA representation baseline. Cells were then maintained at >1,500 cells per guide throughout the screens (see Supplementary methods for more details) and finally harvested at 33-day (day33) post puromycin selection to assess gene essentiality.

### Computational analysis of CRISPRko and CRISPRi data

FASTQ files were analyzed with MAGeCK version 0.5.6 (Li et al., 2014), using RRA and total read count normalization, with default settings. Each replicate was analyzed independently by comparing guide RNA at day33 against day0. CRISPRko and CRISPRi essentiality signatures for each replicate were computed by transforming the p-value to z-score (the sign was inferred from fold change). Lineage and Morphogenic essentiality signatures were computed by integrating the z-scores of the three, same-subtype cell lines, using Stouffer’s method (Stouffer, 1949). Finally, a differential Lineage vs. Morphogenic essentiality signature was computed by comparing the essentiality in Morphogenic vs. Lineage cells. To define a subtype-independent essentiality signature we integrated the essentiality signatures as assessed by CRISPRko and CRISPRi, across all six cell lines, independent of subtype, using Stouffer‘s z-score method (Stouffer, 1949).

### Transcription factor overexpression assay (PLATE-Seq)

Full-length open reading frame (ORF) clones for the top 8 Lineage MRs were ordered from CloneID (Harvard Medical School) and cloned into modified Tet-O-FUW lentiviral expression vector (addgene #30130), which include the puromycin resistance gene. MCherry and EGFP ORFs were used as negative controls in the assay. All clones were sequence verified. For each ORF we introduced a unique 20 bp barcode sequence located 200 bp upstream of the lentiviral 3’-long terminal repeat (LTR) region, as reported in Parekh, U., et al (Parekh et al., 2018). This produces a polyadenylated transcript, which contains the barcode proximal to its 3’ end. All viruses were produced and viral titers were measured individually for each virus. ORF containing lentivirus were transduced into KP4 morphogenic cell lines at MOI = 2 in triplicate (6-well format in the presence of polybrene). In a second triplicates set, lentiviral ORFs were co-transduced with M2rtTA (FUW-M2rtTA, addgene #20342), a tetracycline-inducible transcriptional amplifier, allowing monitoring of MR overexpression at higher and lower levels (Hockemeyer et al., 2008). At 24h following viral transduction, media was changed and puromycin (2.5ug/ml) and doxycycline (0.6ug/ml) were added to the cells, followed by a 5-day incubation period before total RNA was collected by Direct-zol RNA MiniPrep Plus kit (Zymo Research). 69 RNASeq profiles, corresponding to 23 different conditions in triplicate were generated by PLATE-Seq(Bush et al., 2017) using 100ng of total RNA as template in each well.

### Analysis of transcription factor overexpression (Plate-Seq) data

Single-end PLATE-Seq reads were pseudoaligned to the GRCh38 transcriptome (mRNA and ncRNA) and quantified using Kallisto version 0.44.0 (Bray et al., 2016), with sequence-specific bias correction. Transcript-level counts were aggregated by Entrez-IDs and compared between unperturbed Lineage cells (PATU8988S and HPAFII) and unperturbed Morphogenic cells (KP4). The corresponding gene expression signature was transformed into a protein activity signature and compared to the Lineage-Morphogenic protein activity signature (TCGA-ICGC signature) inferred from patient profiles, to ensure conservation of protein activity signatures. This was done by computing a differential gene expression signature between Lineage and Morphogenic unperturbed cell lines using the Student’s T.test, as implemented in the VIPER package (Alvarez et al., 2016) on the normalized gene expression profiles. The metaVIPER approach has been used to transform this differential gene expression signature into a protein activity signature. The conservation between this protein activity signature (inferred by comparing the unperturbed Lineage and Morphogenic cell lines) and the Lineage-Morphogenic protein activity signature computed from the patients was assessed by two-tailed aREA test (Alvarez et al., 2016). Specifically by assessing the conservation of the 50 most differentially activated and 50 most differentially inactivated proteins of the TCGA-ICGC signature in the protein activity signature inferred from the cell lines.

### Evaluation of cell reprogramming in PLATE-Seq overexpression assay

A differential gene expression signature for each experimental condition (perturbation) was computed by comparing the gene expression profiles of perturbed vs. unperturbed (negative control) cells, in the same experimental background. Specifically, cells transduced with mCherry and EGFP, with or without M2rtTA (negative controls), were used as controls for cells ectopically expressing each TFs, with or without M2rtTA, respectively. Each differential gene expression signature was then transformed into a protein activity signature by metaVIPER approach (Ding et al., 2018). The extent of reprogramming induced by ectopic expression of each MR protein was assessed by measuring the enrichment of cell line specific Lineage-Morphogenic Tumor Checkpoint MRs—as computed by comparing Lineage (HPAFII and PATU8988S) and morphogenic (KP4) cell lines— in protein differentially activated following ectopic expression of each MR, with or without M2rtTA expression, by a two-tailed aREA test.

### MR Interaction Network reconstruction

MR interaction networks were assembled by combining transcriptional/post-transcriptional (i.e., VIPER-inferred) interactions, as assessed by Student’s T-test of the differential expression/activity of a target MR (Lineage or Morphogenic) following ectopic expression of a different Lineage MR, using the VST-normalized log_2_ counts of each perturbation, in triplicate, vs. the pool of three mCherry and three EGFP-transduced KP4 cells (controls). Protein-protein interactions were added from the STRING (Szklarczyk et al., 2017) and PREPPi (Zhang et al., 2012) databases. This provides mechanistic insight into how each of the top 8 Lineage MRs regulates the other Lineage and the top 8 Morphogenic MRs, thus supporting their highly modular structure. MR genes (proteins) whose expression (activity) changes were statistically significant (p ≤ 0.05, by one-tailed Student’s T-test) were considered as putative transcriptional (post-transcriptional) targets and included in the network. Molecular interactions from PREPPI and STRING databases were selected based on confidence-score. Specifically, we sorted all the interactions based on their confidence score and reported only the interactions ranking in the top 25%. Network representation was done using Cytoscape (version 3.8.2) (Shannon et al., 2003).

### Pooled TF overexpression assay (scRNASeq)

The same barcoded constructs discussed in the previous section, representing the top 8 Lineage Master Regulator proteins (and mCherry as neg. control), were used in the single cell overexpression assays. ORF viruses were pooled into two viral pools, such that, on average, 2-3 ORFs were randomly transduced into each single cell (MOI = 0.288 / each virus). The M2rtTA construct was added to the second pool to increase the transcriptional ORFs output. KP4 cells were transduced with the two viral pools in 6-well format in the presence of polybrene. Media was changed at 24h post-infection and puromycin (2.5ug/ml) and doxocycline (0.6ug/ml) were added to the cultures. Cells were incubated for a total of 11 days, before trypsinization, addition of Multiseq barcodes(McGinnis et al., 2019) and 10X chromium library preparation and sequencing. Non-transduced (control) KP4, HPAFII and PATU8988S cells were also included at this stage to represent the pre-treatment Morphogenic cell state (KP4), while non-transduced (control) Lineage cell lines (HPAFII and PATU8988S) were used as a proxy for the desired Lineage endpoint state. All cells (normal KP4, HPAFII, PATU8988S and KP4 transduced pools, +/-M2rtTA, were MultiSeq barcoded and mixed prior to the Chromium-run.After the Chromium-run, the cDNA was amplified with 1ul of 2.5uM MultiSeq additive primer added to the cDNA amplification mastermix. After this the material was divided into 3 portions, the whole transcriptome, MultiSeq barcode and the ORF barcode. MultiSeq barcode and ORF barcode portions were amplified with specific primers (MULTI-seq_TruSeq RPIX & MULTI-seq_Universal I5 for MultiSeq and ORF_BC_amplif_oligo_F & ORF_BC_amplif_R for ORF barcode) and spiked in the final NGS library at 1% and 10% total amounts respectively. For the final analysis MultiSeq barcode data was not used (see supplementary methods).

### Single-cells Demultiplexing Analysis

Single-cell BAM files were generated with the cell ranger pipeline (version 3.0.2), using GRCh38 as reference genome. Variant calling was performed with SamTools (Li et al., 2009) and captured in .vcf file, containing the genomic variations of three PDA cell lines processed by RNASeq (HPAFII, KP4 and PATU8988S). Bam files and vcf files were used as input for Demuxlet (Kang et al., 2018) for demultiplexing analysis.

### Protein activity analysis of Pooled TFs overexpression assay (scRNASeq)

Single-cell UMI-counts were filtered based on the previously described QC-metrics and normalized to CPM. A differential gene expression signature was first computed, by comparing single cells from unperturbed Lineage (PATU8988S and HPAFII) and Morphogenic (KP4) cell lines, and then transformed into a differential protein activity signature using metaVIPER, as previously described in the protein activity analysis of Elyada data set. Finally, enrichment of Linage-Morphogenic Tumor Checkpoint proteins in proteins differentially active in this signature was assessed by two-tailed aREA test, assess reprogramming of Morphogenic cells into a Lineage state.

### Cell reprogramming efficiency assessment in pooled TFs overexpression assays (scRNASeq)

scRNASeq profiles representing cells transduced with the same MR or MR combination were considered as independent isogenic-MR sets. To robustly asses reprogramming efficiency, isogenic-MR sets comprising fewer than 30 cells were removed. To assess the reprogramming potential of independent MRs and MR combinations, we first computed differential gene expression signatures by comparing the pool of scRNASeq profiles for each isogenic-MR set to the pool of unperturbed KP4 cells. These signatures were then transformed to differential protein activity signatures using metaVIPER, as previously described.

For each single cell in a specific isogenic-MR set, reprogramming was assessed based on the enrichment of Lineage-Morphogenic Tumor Checkpoint proteins—as assessed from the differential expression signature of unperturbed Lineage (HPAFII and PATU) vs. Morphogenic (KP4) single cells—in proteins differentially active in that cell, by two-tailed aREA test. Perturbed KP4 single cells showing statistically significant enrichment in the Lineage-Morphogenic signature (FDR < 5% by two-tailed aREA test) were considered reprogrammed. Isogenic-MR sets were then sorted based on the fraction of cells assessed as significantly reprogrammed and p-value was assessed using Fisher’s Exact test (two-tailed test), adjusted for multiple hypothesis testing (FDR), by comparing the fraction of reprogrammed cells in the isogenic-MR sets vs. the negative control set (mCherry +/-M2rtTa).

### Western Blots

Full-length ORF constructs for OVOL2 or mCherry (+/-M2RTTA), as described for the overexpression assays, were used for Western Blots. OVOL2 or mCherry (+/-M2RTTA) were lentivirally transduced in KP4 cells, at MOI = 2, in triplicate, followed by puromycin selection and 5 days incubation in the presence of doxycycline (0.6ug/ml) before reprogramming was assessed. Cells were then lysed, total protein levels were measured with BCA Protein Assay Kit (Pierce), and samples were Western Blotted (see supplementary methods for the antibodies)

### Data availability

RNASeq data generated at CUMC from laser capture micro-dissected samples have been deposited on Gene Expression Omnibus data based with the following GEO ID: GSE143584. CRISPR data and RNASeq data related to cell lines, overexpression assay, single-cell PDXLineag model and single-cell overexpression assay have been deposited on Gene Expression omnibus database with the following GEO ID: GSE161369. scRNASeq profiles from the Elyada Set are available at NCBI dbGaP under the accession number phs001840.v1.p1., while scRNASeq profiles from the Chan-Shen-Yue Set are available on the EGA database (EGA ID=EGAD00010001811; sample names: 100070, 91610, 91706, 95092, 96460).

### Code availability

No unique computational code has been generated for this manuscript. All the computational tools have been indicated in the methods or supplementary methods. The genomic instability package used to infer copy number variation (CNV) from single cell gene expression profiles is made available on Bioconductor (Laise and Alvarez, 2022).

## Notes

### Summary of Updates

This revision substantially improves our understanding of the cellular subtypes of PDAC based on the new experiments and the analysis of additional datasets. In particular, we present a biological understanding of the OP cells presented in the original version, which have been renamed MAPK-inactive (M-) in light of their functional association. We now demonstrate the presence of three developmental lineages of malignant cell types, each harboring both M+ and M- substates, and validate their reproducibility across multiple datasets. Finally, we demonstrate the rapid interconversion of cells between the M+ and M- states using single cell lineage tracing. The manuscript is entirely reorganized around the concept of cellular states and all figures are updated accordingly.

